# Concordance between male- and female-specific GWAS results helps define underlying genetic architecture of complex traits

**DOI:** 10.1101/2022.09.28.509932

**Authors:** Anna K Miller, Calvin Pan, Jacquelaine Bartlett, Aldons J Lusis, Dana C Crawford, Scott M Williams, David A Buchner

## Abstract

A better understanding of genetic architecture will help translate genetic data into improved precision-based medicine and clinical care. Towards this end, we explored the use of sex-stratified analyses for several traits in the Hybrid Mouse Diversity Panel (HMDP) and UK Biobank to better determine trait polygenicity and identify contributing loci. This was accomplished by comparing the direction of allelic effects in males and females in sex-stratified analyses under the hypothesis that loci that are not associated with a trait should have equal chances of trending in the same direction of effect. Instead, we found that even for most alleles that do not meet nominal levels of statistical significance, the direction of effect in the two sexes was highly concordant. Results were consistent with hundreds of loci regulating each mouse trait and thousands of loci regulating each human trait, including traits for which no statistically significant loci were identified using conventional approaches. We also found evidence of likely spurious widespread epistasis. Collectively, these findings demonstrate the importance of stratifying by sex to discover novel associating loci, suggest a new method for identifying biologically rather than statistically significant associations, and caution that widespread marginal effects can lead to phantom epistasis.

## Introduction

Complex traits are affected by many genetic variants that together influence their severity, presentation, and prevalence^1,2^. Genome-wide association studies (GWAS) have successfully identified many such variants for a broad range of complex traits. However, for many traits there are important differences between males and females that may reflect underlying differences in their respective genetic architectures. Most association studies combine sexes in their analyses and adjust for sex as a covariate, but this approach can conceal differences that a sex-stratified approach can reveal. The likelihood that such sex differences exist is consistent with the frequent sex differences often seen in disease prevalence, severity, age of onset, or outcomes^3^. For example, the prevalence of autism spectrum disorder is 4 times higher in males than females, mortality rates associated with COVID-19 infection are elevated in males, and females have double the likelihood of developing multiple sclerosis^4–6^. Although factors beyond genetics likely contribute to these gender differences, it is becoming increasingly clear that genetic factors, both autosomal and sex-linked, contribute to the sexually dimorphic effects seen for many complex traits and diseases. A better understanding of the sex differences underlying complex traits and diseases between males and females may improve the accuracy and efficacy of precision medicine-based clinical care as well as reduce health care disparities resulting from the historical relative lack of attention given to women’s health issues.

Although sex is typically accounted for as a variable in sex-combined association studies, a number of studies in both humans and model organisms have successfully improved the identification of genetic variants associated with a range of complex traits and diseases by performing sex-stratified analysis or testing for Genotype x Sex interactions^7–15^. These studies have often identified associations between specific variants or loci and their respective phenotype that were otherwise not identified in a sex-combined analysis or in the analyses of the opposite sex. In one study comparing sex-specific and combined-sex analyses of major depression based on the UK Biobank datasets, sex-specific polygenic risk scores were better at predicting major depressive disorder than sex-combined polygenic risk score, indicating the importance of considering sex in genomic analyses for precision-based medicine going forward^14^. Similar findings were observed with testosterone levels, interactions between polygenic risk scores and sex for a number of blood cell traits and coronary artery disease risk, and cognitive function based on a polygenic risk score for schizophrenia^16–19^. These studies highlight the importance of considering sex as an important variable, and in certain instances performing sex-specific analyses of genetic data to identify variants associated with risk or calculating polygenic risk scores that do not transcend sex.

In addition to the sexually dimorphic loci that are identified in sex-stratified genomic analyses, we reasoned that these studies provide the opportunity for large scale comparisons between the estimated effects of genetic variants on a given trait or disease between males and females. Under the null hypothesis that GWAS variants with a p-value larger than an arbitrary threshold (typically 5×10^-8^ in human studies) have no effect on a phenotype, the expectation is that there should be no relationship between the effects attributed to a particular variant in both independent male and female analyses. However, if there is a real biological association between a genetic variant and a given phenotype, then one would hypothesize that these variants might be enriched for similar effects in separate male and female analyses relative to that expected by chance. Applying this comparison across all genetic variants can provide an additional means of estimating the true number of variants that contribute to trait heritability, independent of the arbitrary p-value threshold for significance that is used in most GWAS studies. Recent proposals to modify p-value thresholds, apply other measures of confidence, or even abandon p-values, highlight the growing sense that p-values alone are not reliable measures of biological significance and should instead be integrated with other information such as prior knowledge, mechanistic support, and data quality^20–22^. Alternative methods of incorporating biological information into statistical thresholds for significance have also been proposed and include measures of biological function such as chromatin accessibility or enhancer status or interactions with promoter regions^23,24^, applying pathway-based approaches as opposed to single variant analyses^25^, machine learning algorithms to interpret GWAS results^26^, and simply relaxing significance thresholds^27,28^.

To examine the role of sex-specific analyses on the ability to detect associations between genotypic and phenotypic variation, as well as test the hypothesis that large-scale comparison between the sex-specific analyses could reveal novel insights into genetic architecture, we compared a series of sex-specific and sex-combined analyses on metabolic and blood-related traits in the Hybrid Mouse Diversity Panel (HMDP). Prior studies have had initial success detecting associations for a wide range of mouse complex traits utilizing the HMDP, a population of more than 100 inbred and recombinant inbred mouse strains designed to measure and identify the genetic and environmental factors underlying complex traits^29^. Compared to GWAS in humans, the HMDP better controls variation due to environmental factors, has many relevant tissues available for phenotyping, and the strains can be used for the integration of follow-up studies. The HMDP also provides an order of magnitude greater mapping resolution compared to traditional linkage analyses in mice^29,30^. As evidence of these advantages, use of the HMDP has successfully identified novel genes for a diverse array of traits, including obesity, diabetes, atherosclerosis, and heart failure, many of which have been validated using transgenic or knockout models^29,31–33^. In addition, previous studies of the HMDP using sex-specific analyses demonstrated an improved ability to identify novel loci related to metabolic disease^8^. Given the outstanding questions related to defining the genetic architecture of complex traits in both males and females, the HMDP represents a powerful tool for identifying specific variants that regulate complex traits in mice, as well as discovering more complex patterns of how genetic variation influences trait variation and disease susceptibility^34^.

In this study, we report a comparison between sex-stratified and sex-combined association analyses of several metabolic and blood traits using the HMDP. The male and female genotype and trait data were published by Bennett et al, but only a sex-stratified analysis of female mice was previously described^35^. The findings in this study demonstrate an improved ability for detecting associations in a sex-stratified analysis in the HMDP. Additionally, they indicate that hundreds of additional loci throughout the genome contribute to the genetic architecture of complex traits, far beyond those identified using conservative and arbitrary p-value cutoffs, even for complex traits for which not even a single statistically significant locus was identified. Because of sex-specific findings for BMI and height in humans using data from the UK Biobank study^36^, we performed sex-stratified analyses in the UK biobank similar to what we did in the HMDP to assess whether patterns of association transcend species.

## Results

### Heritability

To determine the additive and non-additive heritability components for a series of complex traits in the HMDP, narrow sense and broad sense heritability were calculated separately for male and female mice for 8 metabolic traits and 3 blood related traits (Table 1). On average, the metabolic traits had higher heritability than the blood traits in both sexes. Broad sense heritability was greater than narrow sense heritability in males and females for all traits except adiposity in males. The heritability estimates differed between the two sexes for many of the traits. For example, the narrow sense heritability of adiposity was less in females (0.566) than in males (0.740) and the narrow sense heritability of body weight was larger in females (0.598) than males (0.463). However, on average, heritability was similar between the sexes.

**Table 1.** Narrow Sense and Broad Sense Heritability of Metabolic Traits in HMDP.

|  | Females |  | Males |  |
| --- | --- | --- | --- | --- |
| Trait | Narrow Sense | Broad Sense | Narrow Sense | Broad Sense |
| **Metabolic Traits** |  |  |  |  |
| Adiposity | 0.566 | 0.681 | 0.740 | 0.718 |
| Body Weight | 0.598 | 0.691 | 0.463 | 0.598 |
| Cholesterol | 0.400 | 0.438 | 0.257 | 0.352 |
| Fat mass | 0.574 | 0.691 | 0.626 | 0.688 |
| Glucose | 0.175 | 0.290 | 0.122 | 0.211 |
| HDL | 0.140 | 0.206 | 0.144 | 0.211 |
| Insulin | 0.212 | 0.285 | 0.407 | 0.521 |
| Triglycerides | 0.238 | 0.321 | 0.273 | 0.368 |
| **Metabolic Average** | **0.363** | **0.450** | **0.379** | **0.458** |
| **Blood Traits** |  |  |  |  |
| Granulocyte % | 0.262 | 0.358 | 0.239 | 0.298 |
| Monocyte % | 0.199 | 0.325 | 0.121 | 0.208 |
| White Blood Count | 0.118 | 0.205 | 0.044 | 0.101 |
| **Blood Average** | **0.212** | **0.314** | **0.135** | **0.200** |
| **Overall Average** | **0.317** | **0.408** | **0.312** | **0.407** |

### Genetic Correlation of Traits

Given that heritability estimates differed between sexes for some of the traits, sex-stratified genetic correlations were calculated to measure the genetic trait variability within and between sexes. The genetic correlations between pairs of traits were first compared within each sex. The absolute value of the correlation between traits within the same sex was on average ± 0.34 (SD = 0.23) in males and ± 0.29 (SD = 0.21) in females. Related phenotypes were highly correlated in both sexes. For example, cholesterol and triglycerides were highly correlated in both males (0.88) and in females (0.65) and body weight and insulin were highly correlated in both males (0.63) and females (0.76) (Table S1). Some traits were more strongly correlated in one sex than the other. For example, fat mass and insulin were more correlated in females (0.83) than in males (0.43) (Table S1). Additionally, body weight and monocyte % were more correlated in males (0.52) than in females (0.13). Some traits were not correlated such as adiposity and insulin in both males (−0.06) and females (−0.04). Additionally, adiposity and cholesterol were not correlated in either males (−0.02) or females (−0.08).

The genetic correlation of each trait was also calculated between male and female mice. The average of the absolute value of the correlations was 0.58 (SD = 0.22). Some traits had strong correlations across sexes including body weight (0.88) and glucose (0.86) (Table S2). Other traits had much lower correlations across sexes, including HDL (0.17) and white blood count (0.35) (Table S2). While traits were on average more correlated across sexes than between traits of the same sex, some traits were more correlated to other traits within the same sex than to the same trait across sexes. For example, HDL is not very correlated across sexes (0.17) but is much more strongly correlated with triglycerides in males (−0.69) and with glucose in females (−0.46) (Table S2). A similar trend is seen for white blood count where the trait is not very correlated between sexes (0.35) yet is highly correlated with insulin in males (−0.60) and with granulocyte % in females (−0.76) (Table S2).

### Detection of Loci with Main Effects

Given that the heritability estimates and genetic correlations underlying some metabolic traits were variable between males and females, main effects were determined using sex-stratified analyses. Main effect associations for 8 metabolic traits were tested in females using 694 mice encompassing 98 strains from the HMDP. The Bonferroni-corrected threshold for significant main effects, based on the number of haplotype blocks and independent SNPs not within any of the haplotype blocks tested, was p < 4.13E-6. At this threshold, there were no significant marginal effects in females (Table 2). While we were unable to detect statistically significant main effects using a conservative Bonferroni threshold, we identified associations that were previously observed by Bennett et al. in this HMDP dataset^35^. That study used a slightly different strain composition and an FDR-based threshold for significance at p < 1.3E-5^35^. Using the less conservative threshold, we replicated 5 unique SNPs in adiposity and 1 unique SNP in HDL that were discovered as significant in the Bennett et al. analysis (Table S3).

**Table 2.** Detection of Marginal Effects in Female and Male Stratified Mice and in Combined Sexes in the HMDP.

|  | Trait | Female<br>Main Effects | Male<br>Main Effects | Sex-Combined<br>Main Effects |
| --- | --- | --- | --- | --- |
| Metabolic | Adiposity | 0 | 58 | 1 |
|  | Body Weight | 0 | 22 | 6 |
|  | Cholesterol | 0 | 0 | 0 |
|  | Fat Mass | 0 | 60 | 5 |
|  | Glucose | 0 | 0 | 0 |
|  | HDL | 0 | 0 | 0 |
|  | Insulin | 0 | 0 | 0 |
|  | Triglycerides | 0 | 0 | 0 |
| Blood | Granulocyte % | 0 | 0 | 1 |
|  | Monocyte % | 0 | 0 | 0 |
|  | White Blood Count | 0 | 0 | 0 |

While these traits were previously analyzed in female mice by Bennett et al, the same traits were not analyzed in males. Thus, main effects of the same 8 metabolic traits were assessed in males using 664 mice from 102 strains from the HMDP. The Bonferroni-corrected threshold for significant main effects was based on the number of haplotype blocks and SNPs independent of the haplotype blocks tested. The main effect significance threshold was p < 2.89E-6. There were multiple significant marginal associations detected for adiposity (n=58), bodyweight (n=22), and fat mass (n=60), which are the traits with the largest narrow sense heritability among those studied (Table 2, Table S3). We also found an enrichment for main effects over that expected by chance (Table S5). As adiposity, body weight, and fat mass are highly related, we examined whether the significant SNPs overlapped between these traits. Overall, 20 haplotype blocks contained SNPs associated with all three traits. Thirty-five additional haplotype blocks contained SNPs associated with two of the traits, including thirty-three blocks associated with both adiposity and fat mass, 1 block associated with both adiposity and body weight, and 1 block was significantly associated with both body weight and fat mass (Fig. S1, Fig. S2, Table S6). Only 10 haplotype blocks were unique to one of the three traits (Fig. S1, Table S6). Among the loci identified, there was little overlap with a similar analysis of the HMDP mice when fed a different obesogenic diet (Table S7)^8^, suggesting a strong Gene x Environmental component to the sex-specific loci identified in each study.

In addition to studies of metabolic traits, and to test the generalizability of any findings to complex traits more broadly, main effects for 3 blood-related traits were assessed in females using 670 mice encompassing 97 strains and in males using 640 mice encompassing 98 strains, using the HMDP. The Bonferroni-corrected threshold for significant main effects was again based on the number of haplotype blocks and SNPs outside of the haplotype blocks tested. The main effect significance threshold was p < 2.87E-6 in females and p < 2.91E-6 in males. At these thresholds, there were no significant marginal effects detected in either males or females (Table 2).

GWAS often only measure main effects using a sex-combined analysis with an adjustment for sex. Thus, in addition to the sex-stratified analyses, we also analyzed the HMDP data using a combined sex approach to assess if similar or different main effects could be detected. Main effects for the 8 metabolic traits and 3 blood traits were assessed in the sex-combined analysis with both male and female samples, adjusting for sex as a variable in the regression analyses. The Bonferroni-corrected threshold for significant main effects was p < 2.87E-6, and thus similar to that for each of the sex-stratified analyses. At this threshold, there was 1 main effect locus detected for adiposity. However, this region was not one of the 58 loci detected in the male-only analysis. There were also 6 main effect loci detected for body weight, with 4 of these also observed in the male-only analysis. There were 5 main effect loci detected for fat mass, with 4 of these loci also observed in the male-only analysis. There was also 1 main effect locus detected for the blood trait granulocyte % that was not detected in the male-only analysis (Table 2). Thus, in total, of the 13 total marginal effect loci detected in the sex-combined analyses, only 5 were not also detected in the male-specific analysis. Conversely, among the 140 loci detected across all traits in the male-specific analyses, only 8 were detected in the combined sex analyses (Table 2). These results indicate that the power to detect QTLs in the HMDP is diminished in sex-combined analyses that adjust for sex as a variable, rather than conducting separate sex-stratified analyses.

### Normalization of Trait Data Did Not Significantly Alter QTL Detection

Metabolic and blood traits were tested for normality using the Shapiro-Wilk’s method to measure if a non-normal distribution could be driving the male-specific main effects. Nearly every trait in both sexes was non-normally distributed (Table S8). Therefore, to test whether the significant loci identified above were artifacts due to a non-normal distribution of the trait data, phenotypes were normalized using both a rank-based normalization and a Box-Cox normalization. Performing QTL detection for adiposity using the rank-based normalized data resulted in the identification of 54 significant QTLs, whereas using the Box-Cox normalized data resulted in the identification of 63 significant QTLs. All but 1 of the rank-based normalized QTLs were also detected in the unadjusted model (Table S9). There were more Box-Cox significant QTLs detected than were found in the unadjusted model, but all of the QTLs detected based on the unadjusted data were also identified in the Box-Cox normalized data. Thus, there were nearly identical results, regardless of whether the regression analyses were performed on unadjusted or normalized trait data. In addition to these results for adiposity, similar findings were observed for all other traits studied for both the sex-stratified and sex-combined analyses (Table S9). Collectively, this demonstrates that the QTL mapping findings were robust to transformation of the trait data.

### Removal of Outliers Did Not Significantly Alter QTL Detection

In addition to testing whether normalization of the trait data significantly altered the QTL mapping studies, we sought to test whether the QTL detection results were being driven by either a few outlier mice or outlier strains of mice. To test this, outliers were removed from the phenotypic data that were greater than three standard deviations from the mean. Outliers were removed either by removing individual mice that met this criterion or removing strains that collectively met this criterion. When individual mouse outliers were removed from male mice, there were 35 adiposity main effects detected and all 35 of these effects were previously found in the unadjusted male only model (Table S9). For body weight, 24 significant main effects were detected, again, with all these loci already detected in the unadjusted male only model (Table S9). There were 55 significant main effects detected for fat mass with 53 of these effects detected in the unadjusted male only model (Table S9). Of the traits that detected main effects in the unadjusted model, none of the adiposity traits (adiposity, body weight, or fat mass), had outlier strains that were removed due to their average strain values and the criteria described above. Insulin and granulocyte % had strain outliers, but both traits failed to detect QTLs using the unadjusted, normalized, or outlier removed data. Thus, as with data normalization, removal of outliers had only minor effects on the detection of loci that control these metabolic traits in the HMDP.

### Randomization of Sex Prevented Detection of Most Sexually Dimorphic QTLs

After applying various quality control measures to the trait data in the HMDP prior to QTL detection, one of the most striking findings, which was robust to these modifications, was the difference in the number of QTLs that associate with adiposity-related traits in male mice relative to the lack of QTLs identified that associate with adiposity-related traits in female mice. This was surprising, as the overall heritability across all three of the adiposity-traits was similar between males and females, and the traits were highly genetically correlated across sexes (0.55-0.88) (Table S2). Thus, to test the robustness of these findings, the sex of each mouse was randomized as either a male or a female and the linear regression analysis was performed again. Adiposity was analyzed as it had the largest average narrow sense heritability across sexes (0.653) (Table 1). Main effects were detected for adiposity with sex randomly assigned for 100 permutations. Among the 100 permutations, there were few instances where significant main effects were detected in either females (mean = 0.41, SD = 2.04, range= 0,17) or males (mean = 0.79, SD = 2.06, range= 0,13) (Table S10), indicating that the sex-specific QTLs were not likely to be artifacts, but instead represented true sexual-dimorphic QTLs.

### Concordant Directionality of Marginal Effects in Male and Female Mice

Given the increased number of significant main effects detected in males relative to females, we wondered whether genetic associations in females had the same direction of effect on each trait as that detected in males, even if they did not reach the level of statistical significance in females. For example, SNP rs48316748 was the most significant SNP associated with body weight in male mice in the HMDP (p < 1.27E-11, Beta = 2.94), with the “A” allele being associated with increased body weight. The “A” allele at this SNP was also associated with an increase in body weight in the analysis of body weight in females, although the p-value was not statistically significant (p = 0.11, Beta = 0.74) (Fig. S3). To test the concordance of direction of effect among statistically significant loci, the direction of effect of each SNP was assessed in each sex and compared. If there were no effects shared between male and female mice, the null hypothesis is that the direction of effects in the two sexes should be the same ∼50% of the time, based on the equivalence of random effects. However, among two of the three traits for which significant main effects were detected in males (body weight and fat mass), the allelic direction of effect was the same in males and females significantly more frequently than by chance alone (Table 3). The enrichment in the number of significant main effects with the same allelic direction of effect was significant in body weight (p = 4.20E-8) and fat mass (p = 0.003) but not in adiposity (p = 0.22) (Table 3). Therefore, although the significant main effects may not be statistically significant in both sexes, they nonetheless had a consistent effect direction in both male and female mice (Table S11).

**Table 3.**
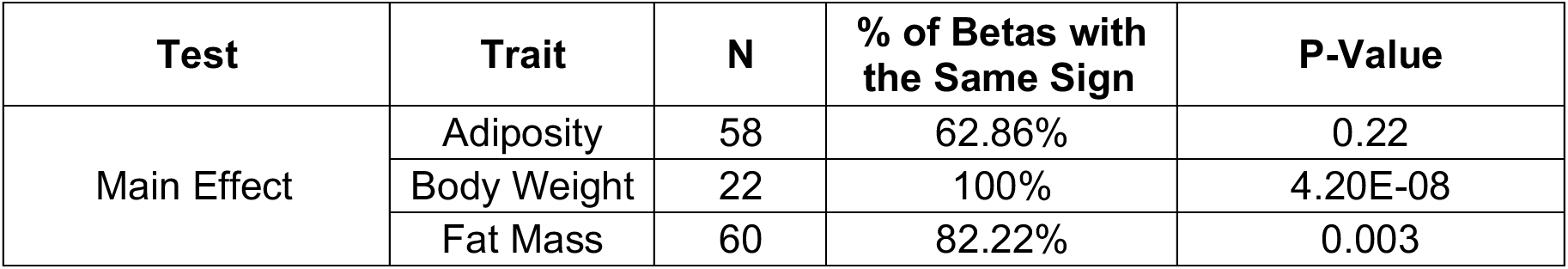
Bonferroni Significant Main Effect Compared Between Sexes.

| Test | Trait | N | % of Betas with the Same Sign | P-Value |
| --- | --- | --- | --- | --- |
| Main Effect | Adiposity | 58 | 62.86% | 0.22 |
|  | Body Weight | 22 | 100% | 4.20E-08 |
|  | Fat Mass | 60 | 82.22% | 0.003 |

To extend these findings beyond the loci with p-values that met the conservative Bonferroni-adjusted threshold for significance, we next tested whether the direction of effects was similar in males and females for all main effects tested, regardless of statistical significance. Concordance between males and females were analyzed and sorted based on the p-value for association in males. Male p-values were used as all significant main effects were detected in males. For all traits examined, the percentage of loci with effects in the same direction in males and females was far greater than the 50% expected by chance (Fig. 1A, Fig 1B) across nearly all levels of statistical significance. The enrichment relative to chance was greatest among the most significant loci and decreased towards the 50% expected by chance as the male p-values increased, asymptotically approaching 50% as p-values approached 1.

**Figure 1.**
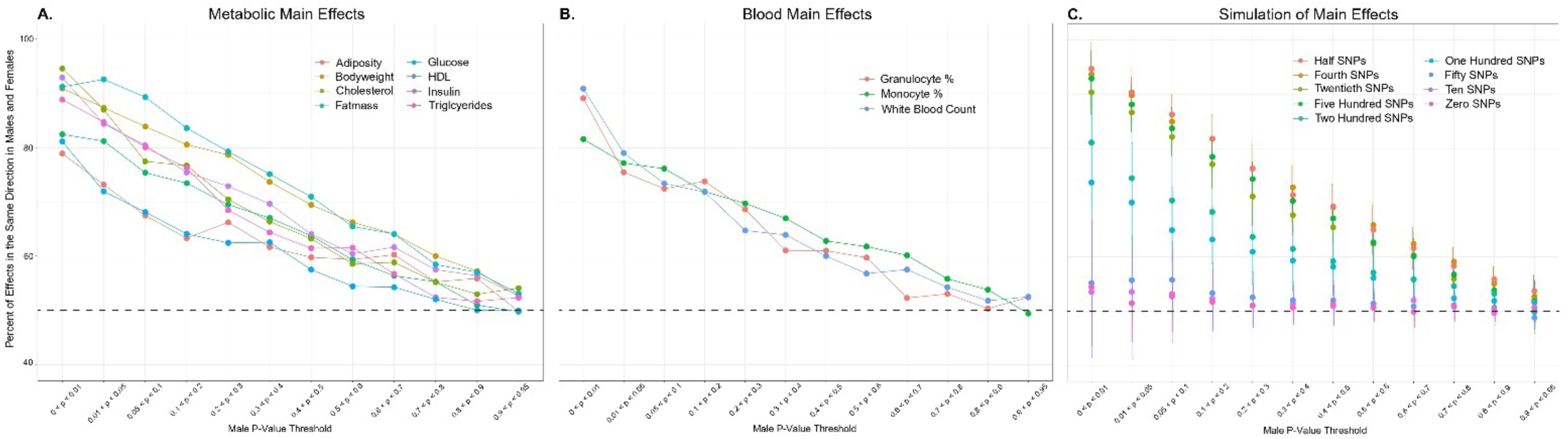
Detection of Widespread Main Effects in the HMDP. The percentage of all HMDP main effect loci with consistent direction of effects in the two sexes were plotted according to the male derived p-values. Main effect p-values > 0.95 were excluded due to many low frequency alleles. Directionality was determined using the sign of the effect size of each main effect loci. The phenotypes measured included (A) metabolic traits, (B) blood traits, and (C) simulated phenotypes using an increasing number of “causal” SNPs.

To estimate the number of SNPs with heritable effects on a given trait that would be consistent with the observed enrichment of greater than 50% of loci with effects in the same direction in males and females, phenotypes were simulated based on the exact HMDP genotypes. Phenotypes were simulated with a heritability of 0.4 to approximate the heritability of the metabolic traits (Table 1). To recapitulate the independent analysis of male and female mice, all SNPs in the simulation were assigned an artificial beta value. For SNPs defined as having a “true” causal effect that was consistent in both male and female mice, these SNPs were assigned beta values in the same direction (i.e. both positive or both negative) in both the male and female simulation. To simulate a SNP with no causal effect, beta values were randomly assigned separately in the male and female analysis and were thus on average in the same direction 50% of the time. Phenotypes were simulated with a range of “causal” SNPs, from a minimum of zero causal SNPs to a maximum where half of the total number of SNPs are estimated as causal (n=10,998). After independently simulating the phenotype data under each of these conditions 10 times, linear regression analyses were performed on each of the 10 simulations as was done for the actual trait data.

As expected, in the absence of any “causal” SNPs in the simulation, there was no enrichment of male and female concordance above the 50% expected by chance (Fig. 1C). In contrast, when at least 100 “causal” SNPs were included in the model, a clear increase in the percentage of concordant effects in males and females was detected relative to the 50% expected by chance (Fig. 1C). Comparing the results from the real phenotype data to the results from the simulated phenotypic data, the metabolic and blood traits likely include between 200 and 1,100 SNPs with a true marginal effect (Fig. 1). These data demonstrate that the marginal effects detected for many loci throughout the genome, even though they do not reach the conservative p-value threshold for significance, are not simply random noise and instead indicate that these loci are in fact contributing to trait heritability via their marginal effects.

### Phantom Detection of Loci with Interaction Effects

Given the increased power of using male/female concordance to estimate the number of contributing loci compared to a conservative Bonferroni threshold for significance, we wanted to assess whether this could be applied to SNP-SNP interactions, as the role of epistasis in complex traits remains controversial^37–39^. Replicating the methods described for main effects, we tested whether the direction of effects was similar in males and females for all pairwise interactions analyzed, regardless of statistical significance. Concordance between males and females were measured and sorted based on the p-value for association in males. For all traits examined, the percentage of loci with effects in the same direction in males and females far exceeded the ∼50% expected by chance across nearly all levels of statistical significance (Fig. 2A, Fig. 2B). As seen with the main effects, enrichment of concordant interactions relative to chance was greatest among the most significant interactions and decreased towards the 50% expected by chance as the p-values increased, asymptotically approaching 50% as p-values approached 1. To evaluate the number of genome-wide pairwise interactions consistent with these findings, we again turned to the simulations of phenotypic data. Given the estimate that at least 200 main effects contribute to the heritable regulation of each trait, we applied that model of genetic architecture to the detection of genetic interactions. For comparison, we also modeled the effects of zero main effects as a negative control, and >10,000 main effects as a likely overestimation. Importantly, the genetic architecture simulated by the model included only main effects. Thus, if given this simulated genetic architecture no interactions were detected, it would support the validity of widespread interactions as indicated by the elevated concordance of the male/female direction of effects. However, if the simulated genetic architecture, including only marginal effects, recapitulated the elevated concordance of the male/female direction of interaction effects, it would indicate that these findings were artifacts of the widespread main effects and did not represent true interactions.

**Figure 2.**
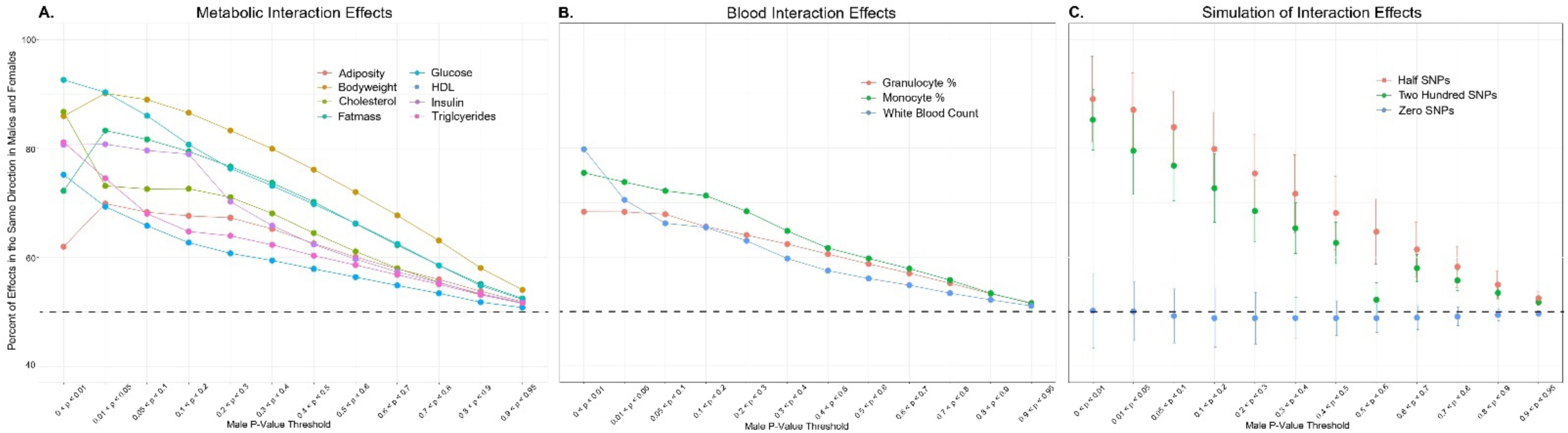
No Widespread Detection of Pairwise Epistasic Interactions in the HMDP. The percentage of all HMDP pairwise interacting loci with consistent direction of effects in the two sexes were plotted according to the male derived p-values. Interaction p-values > 0.95 were excluded due to many low frequency alleles. Directionality was determined using the sign of the Beta value for each interaction. The phenotypes measured included (A) metabolic traits, (B) blood traits, and (C) simulated phenotypes using an increasing number of “causal” SNPs.

As expected, when zero marginal effects were included in the model, there was no overrepresentation of male/female concordance for interactions relative to chance. However, when 200 main effects were included in the model, the male/female concordance rate was far greater than the 50% expected by chance, mirroring the results from the actual HMDP data (Fig. 2C). As the simulated data including only marginal effects had a pattern similar to the actual data, it is consistent with an interpretation that the interaction results were artifacts due to the widespread underlying main effects. There may nonetheless be some epistatic interactions contributing to the phenotype, as the broad sense heritability was higher than the narrow sense heritability, but we were unable to detect the signature of widespread interactions analogous to that which was detected for the marginal effects.

### Concordant Directionality of Marginal Effects in Humans

To test whether the increased frequency of condordant directionality was also observed in human data, as it was in the HMDP (Fig. 1), we performed an analogous comparison of both BMI and height in 219,777 males and 260,528 females using data from the UK Biobank study^36^. As was detected in the HMDP, concordance rates between males and females were again significantly elevated relative to that expected by chance, with the highest concordance rates again observed among the most statistically significant loci, and with levels again asymptotically approaching 50% as the p-values approached 1 (Fig. 3A). We also performed simulation studies to estimate the likely number of loci contributing to the regulation of BMI based on these concordance rates. The negative control experiment, with no simulated main effect loci, as expected demonstrated no enrichment of concordance at any p-value (Fig. 3B). The simulation that most closely matched the actual BMI UK Biobank data was the one where one-third of all tested loci contained a main effect (Fig. 3B), consistent with tens of thousands of loci across the genome contributing to the regulation of this trait. This number of contributing loci with main effects is significantly higher than that estimated from the HMDP mouse data (Fig. 2B), likely due to the increased power obtained using hundreds of thousands of individuals compared to just hundreds of mice, although species differences in genetic architecture cannot be ruled out. Nonetheless, these analyses indicate that an order of magnitude more loci are contributing to the regulation of BMI and other complex traits relative to those identified as statistically significant using similar sized populations^40^ and is instead more consistent with a genetic architecture estimated by GWAS using more than 10x the number of samples^41^.

**Figure 3.**
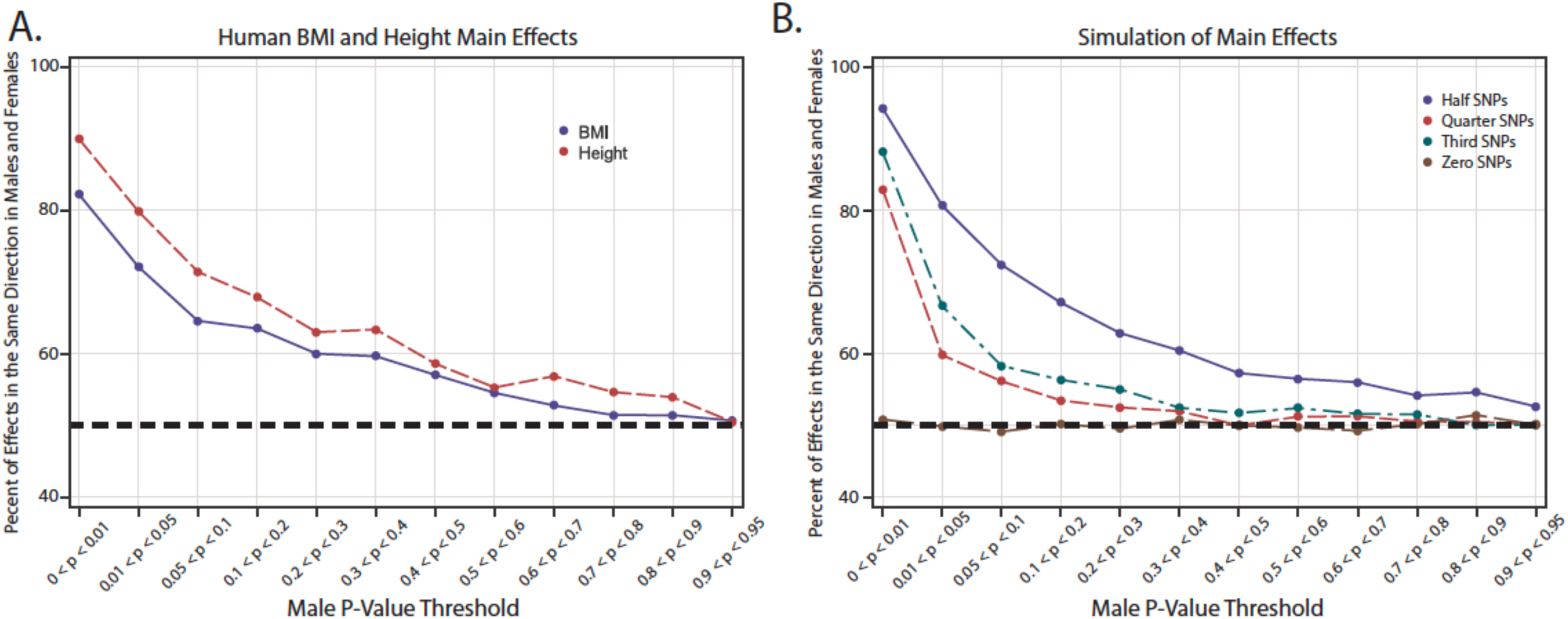
Detection of Widespread Main Effects in the UK Biobank. The percentage of all UK Biobank main effect loci with consistent direction of effects in the two sexes were plotted according to the male derived p-values. Main effect p-values > 0.95 were excluded due to many low frequency alleles. Directionality was determined using the sign of the effect size of each main effect loci. The phenotypes measured included (A) BMI and height (B) and simulated phenotypes using an increasing number of “causal” SNPs.

## Discussion

In the current study, we utilized the HMDP to measure the sex-specific heritability of several metabolic and blood traits. As expected, broad sense heritability was higher than narrow sense heritability across almost all traits and in both sexes, indicating a contribution from dominance or non-additive factors to total heritability. However, the additive component always comprised a majority of total heritability. The contribution of non-additive or dominance factors to total heritability is consistent with many prior studies, as is the smaller contribution relative to additive effects^35,42–45^. It is worth noting that adiposity, body weight, and fat mass, the traits for which significant main effects were identified in males, exhibited the highest levels of narrow sense heritability relative to all other traits. As main effects were detected in males only for these three traits, it is unsurprising that the narrow sense heritability was greater in males than in females for adiposity and fat mass. Surprisingly, narrow sense was greater in females than in males for body weight. This indicated there could be genetic differences underlying body weight that differs by sex.

The underlying genetic differences of metabolic and blood traits were further studied through the genetic correlation of traits across sexes and between traits within a single sex. Body weight was highly correlated across sexes, but adiposity and fat mass were less strongly correlated. Interestingly, some traits were more strongly correlated with another trait within the same sex than to the same trait in the other sex. Thus, the genetic makeup of some traits may be more closely related to other phenotypically related traits in the same sex. We therefore speculate that there are substantial genetic differences regulating some traits across sexes.

This study aimed to better understand the pervasive role of sex-dependent marginal effects in the underlying genetic architecture. Studies were performed using standard linear regression analyses and simulated phenotypes to identify significant marginal associations and non-additive epistatic interactions, and many such loci were discovered (Table 2). Normalization, randomization, and simulation studies all supported the finding of many more marginal associations in males relative to females. However, the simulated phenotype analyses also indicated that detected epistatic interactions were likely the result of spurious signals from marginal effects mimicking an interaction, as has been previously reported^46–48^. Nonetheless, the higher broad sense heritability estimates relative to narrow sense are still consistent with a non-additive contribution, albeit given the statistical difficulties in detecting interactions, functional studies will be needed to discover such interaction effects.

Many detected main effects were sex-dependent as they were detected in analyses of male mice only but were undetected in females alone or sex-combined analyses. Prior human GWAS have detected sex-dependent marginal effects for depression phenotypes^14^, apical periodontitis^49^, Behçet’s disease^50^, and blood pressure^15^. The sex-dependent nature of the significant main effects in our studies was further supported by randomizing sex in the analyses, which abolished the associations, providing an important control for spurious findings in the sex-specific analyses. Nonetheless, the significant main effects in males largely had the same direction of effect in females. This pattern was seen beyond the statistically significant loci, and thus numerous possible small but likely real marginal effects were detected for each trait. The detection of many marginal effects likely contributing to the genetic architecture echoes previous findings of widespread genetic effects^2^, although a different statistical method was used, different traits were examined, and a different organism was studied. Thus, it is perhaps not surprising that analysis of BMI and height in humans demonstrated similar findings as the complex traits analyzed in the HMDP. Together the results support the robustness of a highly polygenic model as a key feature in the genetic architecture of most, if not all, complex traits. It is expected that if statistical power were further increased in the HMDP with the inclusion of additional strains and replicates, to more closely approximate the UK Biobank studies, the number of detected loci may yet be higher, potentially approaching omnigenic inheritance.

Accounting for the highly polygenic, or potentially omnigenic, nature of the genetic architecture may be important for improving the modeling of trait heritability in complex traits driven by many genetic associations. The incorporation of many loci may improve the ability to predict disease risk via more accurate polygenic risk scores (PRS) that account for variability accounted for by sex. As argued previously, we suggest PRS can be used to efficiently model the complex traits using many main effects^51^. As we detected between two hundred and ∼1,000 SNPs to have a marginal effect in mice and many thousands more in humans, PRS could be constructed using the top main effects as estimated based on male/female concordance as an additional method, beyond using an arbitrary p-value cutoff. In the HMDP study, we were able to detect these polygenic effects with only 700 mice from each sex. Thus, this method may be successful in studies of all sample sizes, including studies with reduced statistical power. Using the top main effects that fall within the number of variants suggested to contribute based on male/female concordance rates is an additional method that could more accurately model the contribution of marginal effects in complex traits regardless of sample size.

In conclusion, we present a simple but novel analysis applied to the HMDP and UK Biobank to detect poly- and perhaps omnigenic marginal effects regulating several correlated and uncorrelated complex traits, even for those traits that lacked any statistically significant marginal effects. This indicates that many loci spanning the genome are likely contributing to trait heritability, and strongly argues that current conservative p-value thresholds may not be the best means to approach association studies. Integration of this polygenic model may lead to more accurate genetic models for most complex traits, for which current models fail to account for most heritability and similarly fail to accurately predict trait outcomes based on polygenic risk scores in distinct and diverse populations^42,43,45,52–54^. The results also indicate that there are likely multiple sex-dependent genetic effects/architecture that sex-combined analyses, that even adjusting for sex cannot detect. Therefore, future studies may more effectively model a trait’s heritability and complex genetic architecture by performing sex-dependent analyses as well as including the many main effects that play a role in heritability but are not statistically significant. With a more comprehensive understanding of the many main effects required to model genetic architecture, a clearer understanding of disease etiology will hopefully be detected for many complex traits. Continued advances towards a better understanding of the genetic architecture of complex traits and diseases promises to improve disease risk prediction, help unravel disease pathophysiology, and guide the use of personalized medications and treatments with improved precision.

## Methods

### Mice

The HMDP genotype and trait data are from Bennett et al. and included 8 metabolic traits (adiposity, bodyweight, cholesterol (unspecified), fat mass, glucose (unspecified), HDL, insulin, and triglycerides) and 3 blood traits (granulocyte %, monocyte %, and white blood count) and were obtained from the Mouse Phenome Database (https://phenome.jax.org/)^35,55^. These traits were selected for analysis because they had the largest sample sizes available among all HMDP datasets. Males and females were analyzed both separately (Fig. S4) and in a sex-combined analysis. A total of 694 female mice from 98 strains and 664 male mice from 102 strains were used in the metabolism trait analyses. Strain BXD24/TyJ-Cep290<rd16>/J was removed from both male and female analysis as it carries a mutation that affects vision^56^ and four additional strains were removed due to missing phenotype or genotype data (Table S12). The female analyses also excluded strain BXD29-Tlr4<lps-2J>/J and the male analyses excluded strains AXB13/PgnJ and NON/ShiLtJ due to missing phenotype or genotype data (Table S12)^35^. The blood trait analyses used 97 strains and 670 female mice and 98 strains and 640 male mice. Both the female blood trait and metabolism trait analyses used the same strains. The male blood trait strains were the same as the male metabolism strains except for 3 fewer strains (BXD31/TyJ, BXD6/TyJ, and CXB8/HiAJ were excluded) (Table S12).

### Genotype analysis

The HMDP strains were previously genotyped using the Mouse Diversity Genotyping Array^55^. The number of genotyped SNPs in females was 200,885 and in males was 199,910 (Table S13)^35^. These genotype data excluded multiallelic SNPs; thus, only biallelic SNPs were included in these numbers and used in the analysis. SNPs were further pruned from the HMDP data using the PLINK linkage disequilibrium (LD)-based SNP pruning command using an R^2^ of 0.8^57^ (Table S13). Haplotype blocks were calculated in PLINK from the remaining SNPs using a max block size of 15,000 kb^57^. For the female metabolic traits, 12,221 haplotype blocks plus 2,314 SNPs located outside of any blocks were used in the analysis. For the male metabolic traits, 11,385 haplotype blocks plus 5,907 SNPs located outside of blocks were used in the analysis (Table S3). For the female blood traits, 12,193 haplotype blocks plus 5,221 SNPs located outside of blocks were used for analyses. For the male blood traits, 11,437 haplotype blocks plus 5,752 SNPs located outside of blocks were used in the analysis (Table S13). The total number of haplotype blocks and SNPs for each trait were used to calculate the Bonferroni significance thresholds for female metabolism main effects (3.44E-6), male metabolism main effects (2.89E-6), female blood main effects (2.87E-6), and male blood main effects (2.91E-6). When determining significant marginal associations within a haplotype block, only the SNP with the most significant p-value within the block was used. Similarly, when estimating the direction of effect for pairwise interactions between haplotype blocks or SNPs outside of the haplotype blocks, only the most significant SNP-SNP interactions within a block were considered. The thresholds for significance applied to main effects varied between this analysis and that of Bennett et al., due to the different number of strains analyzed and the FDR-based threshold previously used as opposed to the Bonferroni-based threshold in this study^35^.

In the sex-combined analyses, SNPs were removed that were not present in both male and female genotype data (1,773) or had a differing minor allele (3,922). After removing these SNPs, LD pruning was performed as described above. The sex-combined metabolic and blood traits had 22,284 SNPs remaining resulting in 12,229 haplotype blocks with 5,129 SNPs located outside of the blocks (Table S13). The total number of haplotype blocks and SNPs outside of the blocks were used to calculate the Bonferroni significant threshold for combined main effects (2.87E-6).

### Statistical Analysis (HMDP)

Trait heritability was estimated using two approaches, “narrow sense heritability” and “broad sense heritability”^58^. Narrow sense heritability, which refers to heritability due to additive genetic variance, was calculated using the Genome-wide Complex Trait Analysis (GCTA) software^59^. It estimates the proportion of phenotypic variance explained by all genome-wide SNPs for a complex trait and the narrow sense heritability estimate is based on sharing genomic regions identical by descent^59^. Broad sense heritability, which includes not only additive effects but dominance and other non-additive effects as well^59^, was calculated using the R package “heritability” and was estimated via reproducibility of trait measures in individuals of the same strain^60^.

The genetic correlation for trait data between and within sexes was estimated using GRM and GREML in the Genome-wide Complex Trait Analysis (GCTA) program^59^. GRM estimates the genetic relatedness from SNPs and GREML estimates the variance explained by the SNPs in the GRM^59^. The reml-bivar command was used to compare the variance explained by the SNPs in the GRM for each trait across sexes. It also compared across traits within the same sex, comparing two traits at a time.

FaST-LMM (version 0.6.1), a linear mixed model method that accounts for population structure, was used for single locus main effects and pairwise interaction association tests^61^. A kinship matrix was constructed for the SNPs on all chromosomes except the one being tested to improve power and allows main effect SNPs to be tested only once for association in the regression equation. Not every main effect and interaction was measured in both sexes as differences in haplotype block structure excluded different SNPs. Both interchromosomal and intrachromosomal SNP-SNP interactions were calculated^61^. FaST-LMM calculated the effect size for each SNP as the fixed-effect weight for main effects and as the Beta for each SNP-SNP interaction for epistasis tests^61^. The enrichment of associations identified at p-value < 0.05 were calculated using chi-square tests.

The normality of all phenotypes was measured using the Shapiro-Wilk’s method via the R package “stats” (version 3.6.2). All phenotypes were rank normalized using the R package “RNOmni” (version 1.0.0). Trait data was also Box-Cox normalized using the R package “bestNormalize” (version 1.8.3)^62^. Outliers were removed using two metrics: removing individual mice outside of 3 standard deviations away from the mean and strain averages outside of 3 standard deviations away from the mean.

GCTA was also used to simulate the phenotypic data^59^. The simulated phenotypes were calculated with a heritability of 0.4, similar to the heritability of the metabolic trait data for male and female mice (Table 1). Each simulation was performed separately for males and females to mimic the sex-specific analyses performed with the real HMDP data. Within the genotype data, we modeled SNPs as causal or having no effect. Beta values were assigned to both categories of SNPs. Causal SNPs were assigned a beta value that was identical in both size and direction in the independent male and female simulations. SNPs modeled to have no effect were randomly assigned a normally distributed beta value in each of the male and female simulations, with no link between these separate analyses as to the direction or magnitude of the effect for a given SNP. To mirror the effect sizes detected in the real trait data, the SNPs considered to have no effect were assigned an average beta value of ±0.2, while the SNPs considered to have a causal effect were assigned an average beta value of ±3.0. The number of causal SNPs that were simulated ranged from zero to half of all SNPs and included the following sets of causal SNPs: half, one-fourth, one-eighth, one twentieth, two hundred, one hundred, fifty, ten, and zero. For the simulations where zero SNPs were considered causal, no beta values were included in the simulation so that they were again randomly assigned in each of the male and female simulations. Interactions were tested using simulated data based on either half, two hundred, or zero SNPs considered as causal for a marginal effect, with no true interactions modeled in the simulated data. Each phenotype simulation and analysis of main effects and interactions was repeated 10 times.

Sex-stratified main effect analyses were also analyzed with sex randomized, using 100 permutations. In each permutation, the number of males and the number of females were consistent in each analysis as there were more female mice than males. Randomization was performed in R using the transform command in base R (version 3.6.2).

### Statistical Analysis (UK Biobank)

UK biobank genetic data as well as height and BMI clinical trait data were obtained from the UK Biobank data repository. Trait data was rank normalized in R. Quality control was run on males and females separately consisting of removing non-biallelic variants, LD greater than 0.8, individuals with missing genotype data greater than 2%, genotype missing data greater than 5%, Hardy-Weinberg Equilibrium p-values less than 10E-5, and variants present in both males and females. The complete dataset for BMI and height consisted of 219,777 males and 260,528 females.

FAST-LMM was used to detect marginal effects. To limit computational resources, every 10^th^ variant was included in the analysis for a total of 12,866 SNPs. GCTA was used to simulate the phenotypic data^59^. The simulated phenotypes were calculated with a trait heritability of 0.248, based on the heritability estimates for BMI obtained from the website http://www.nealelab.is/uk-biobank/. SNPs were modeled as either causal or having no effect using the same procedure as described for the HMDP data. SNPs classified as causal were randomly assigned a normally distributed average beta value of ± 0.0235, which was identical in both size and direction in both the male and female simulations. SNPs classified as no effect were randomly assigned a normally distributed average beta value of ± 0.0071 in each of the male and female simulations, with no link to the direction or magnitude between the sexes. The proportion of causal SNPs simulated included one-half, one-third, one-quarter, and zero.

## Data Availability

All genetic and phenotypic data for the HMDP was obtained from the Mouse Phenome Database (https://phenome.jax.org/). All genetic and phenotypic data for the UK Biobank was obtained from the UK Biobank data repository (https://ukbiobank.ac.uk).

## Acknowledgements

This work made use of the High-Performance Computing Resource in the Core Facility for Advanced Research Computing at Case Western Reserve University. We acknowledge Dr. Carl Kadie for his assistance with analyses in FaST-LMM. This research was funded by grants DK119305, NLM010098, and 5T32HL007567-35.

## Author Contributions

A.K.M. led the analysis of the HMDP data and writing of the manuscript. C.P. assisted with analyses of the HMDP data. J.B. led the analysis of the UK Biobank data. A.J.L. provided guidance with the HMDP data analysis. D.C.C. provided guidance for all analyses and assisted with analysis of the UK Biobank data. S.M.W. and D.A.B. together developed the concept and oversaw all experimental design and data analysis. All authors edited and provided feedback for the manuscript.

## Competing Interests

The authors have no competing interests to disclose.

## Materials and Correspondence

Correspondence to David A. Buchner, and Scott M. Williams

## Notes

### Competing Interest Statement

The authors have declared no competing interest.

### Summary of Updates

This version of the manuscript contains many significant changes to the data analysis and conclusions.

