## Supplementary figures and images for "Concordance between male- and female-specific GWAS results helps define underlying genetic architecture of complex traits"

### Figure S1

#### Adiposity

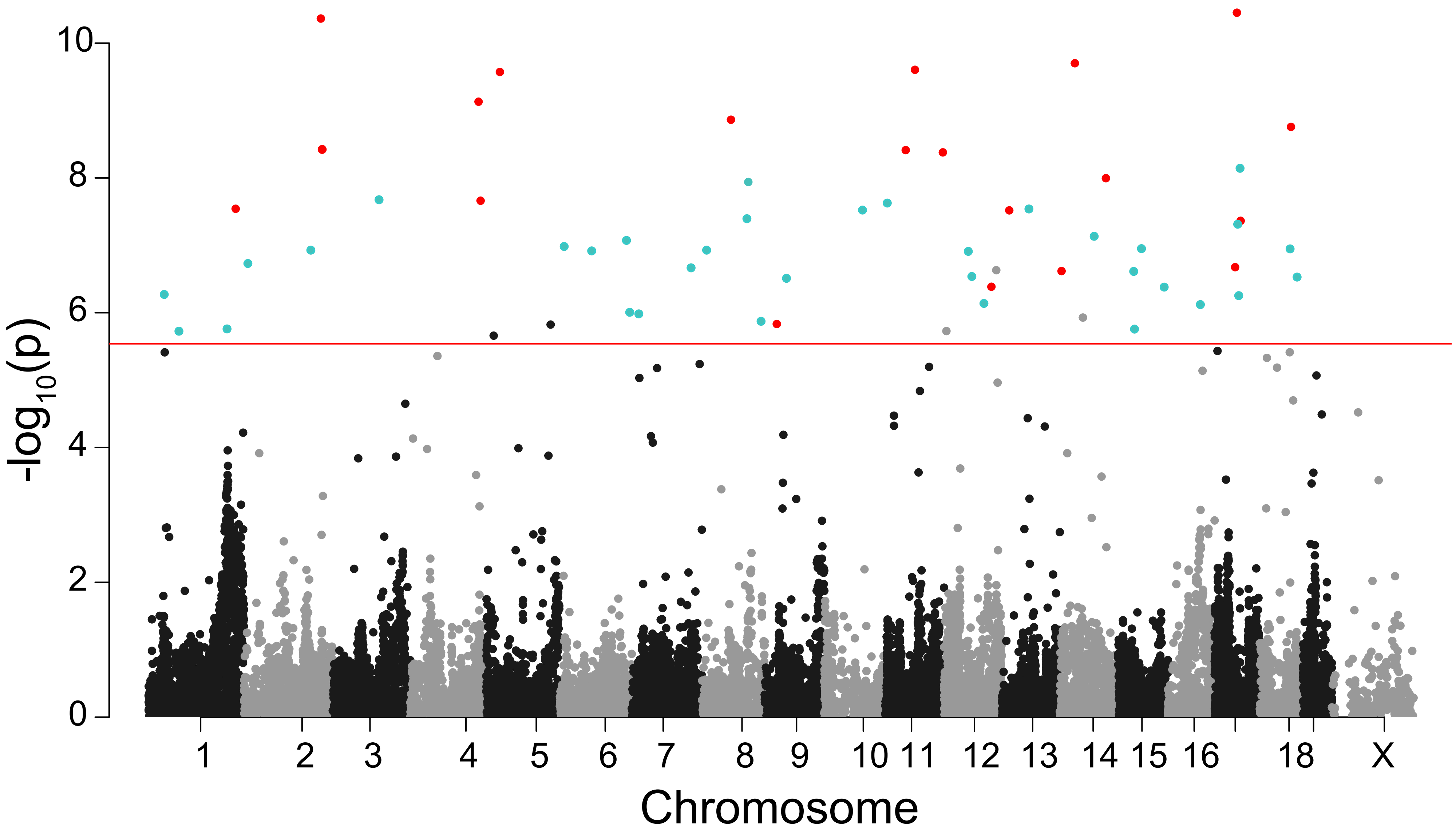


#### Body Weight

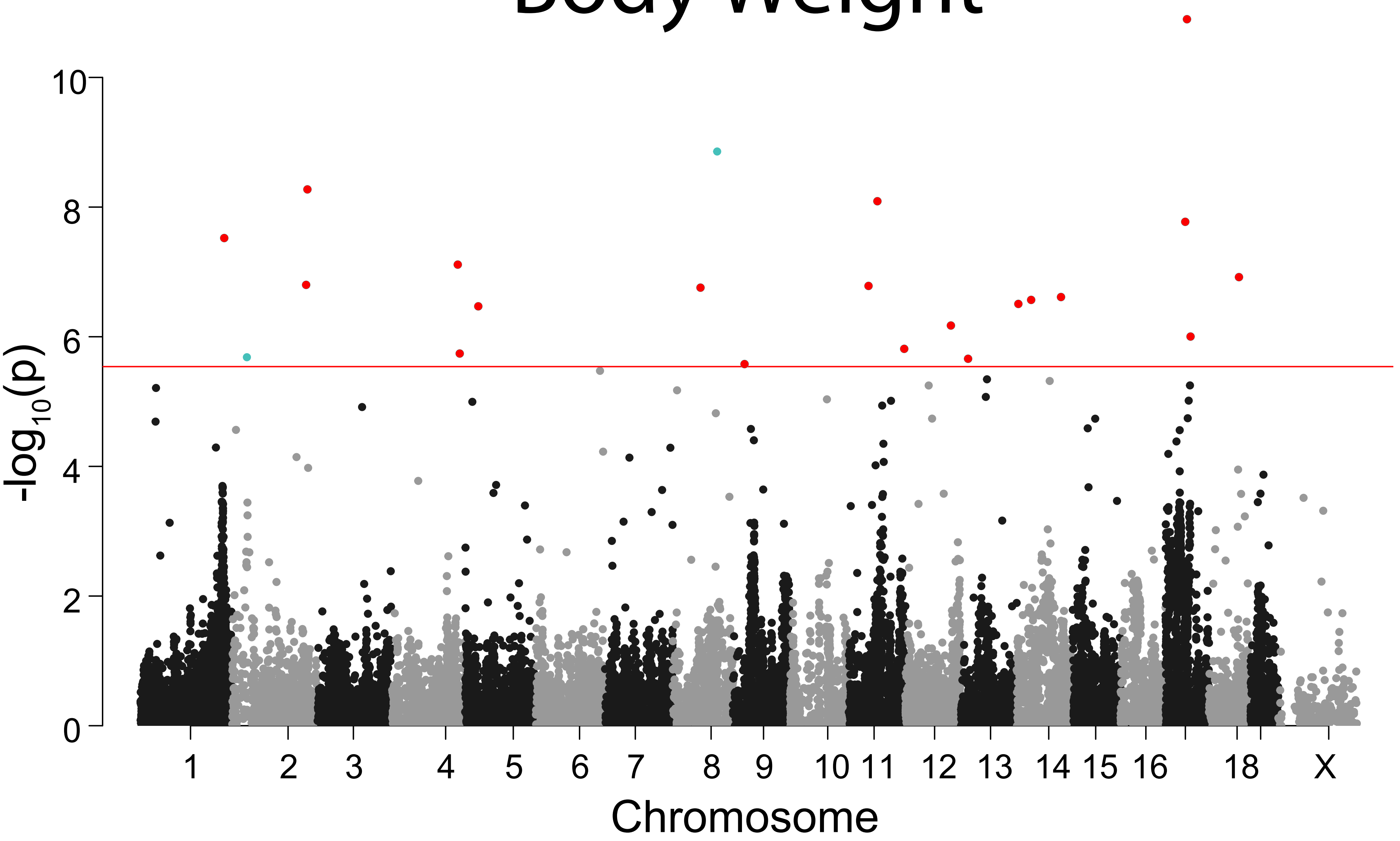


#### Fat Mass

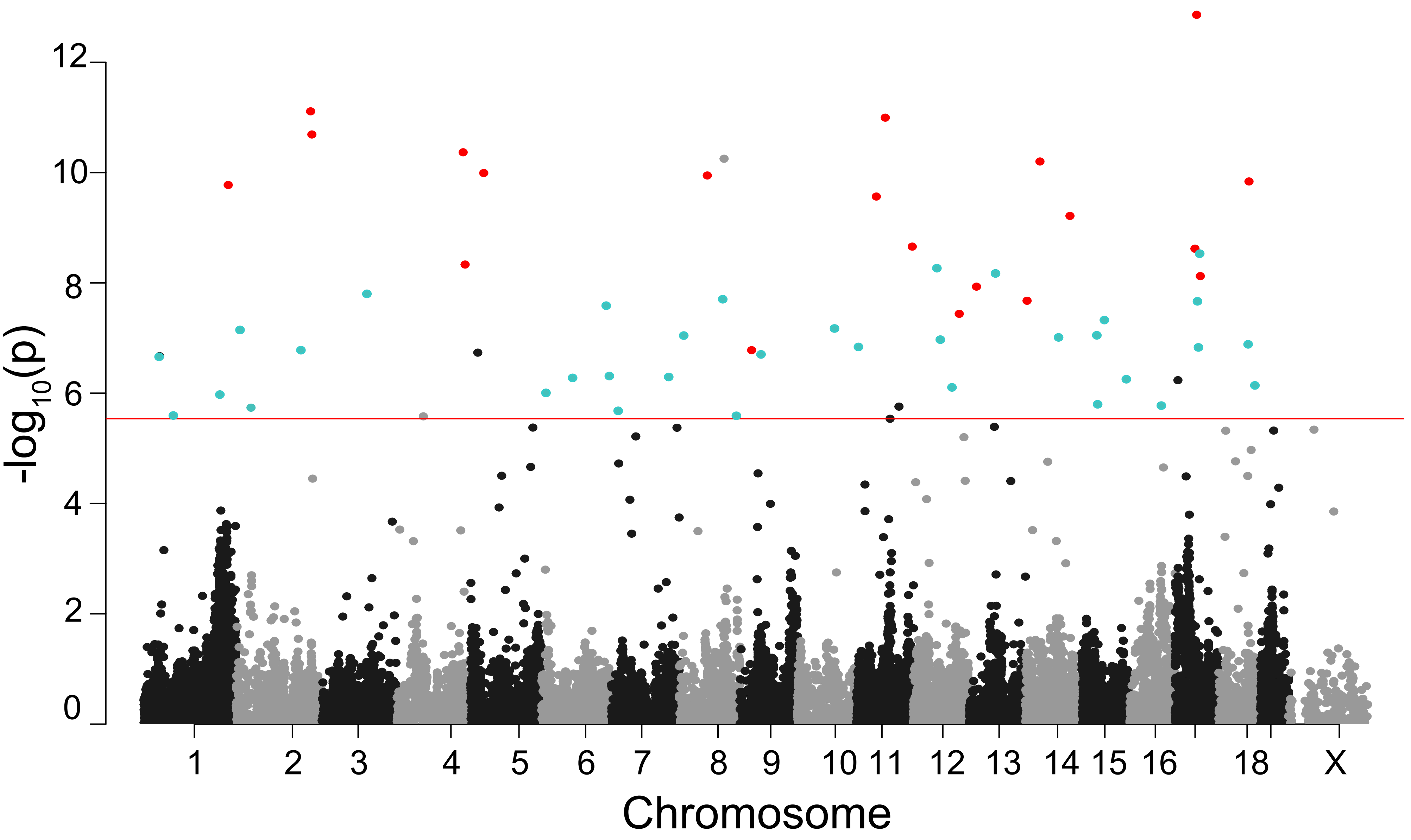

### Figure S2

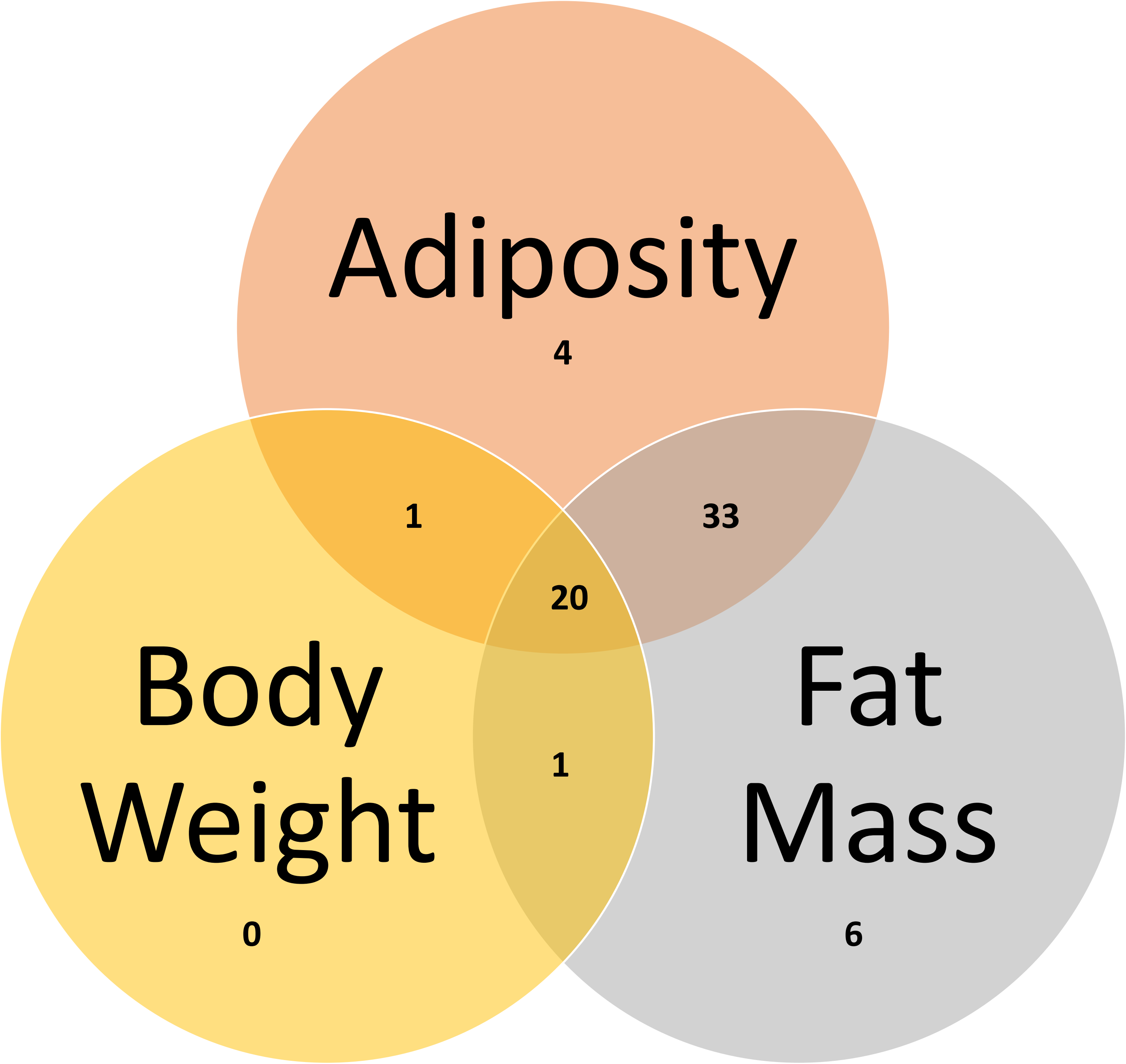

### Figure S3

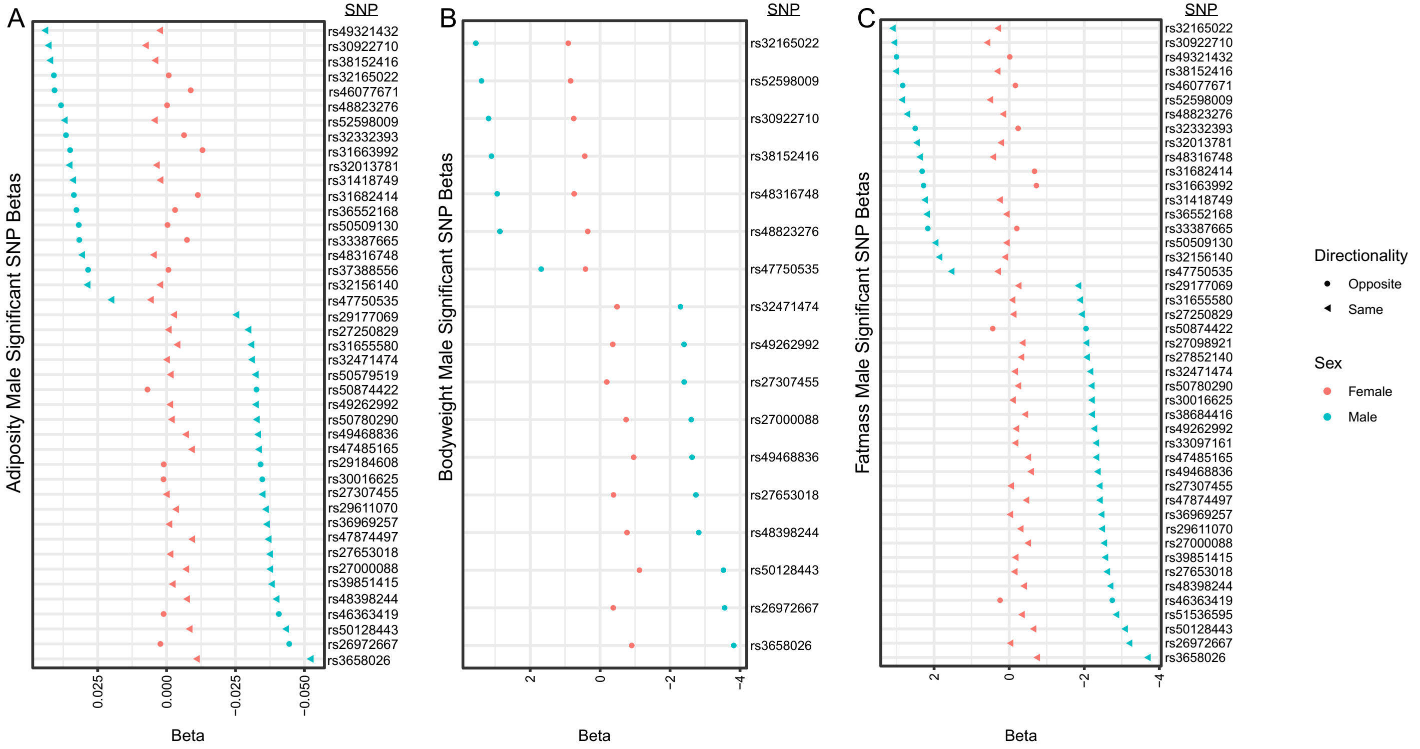

### Figure S4

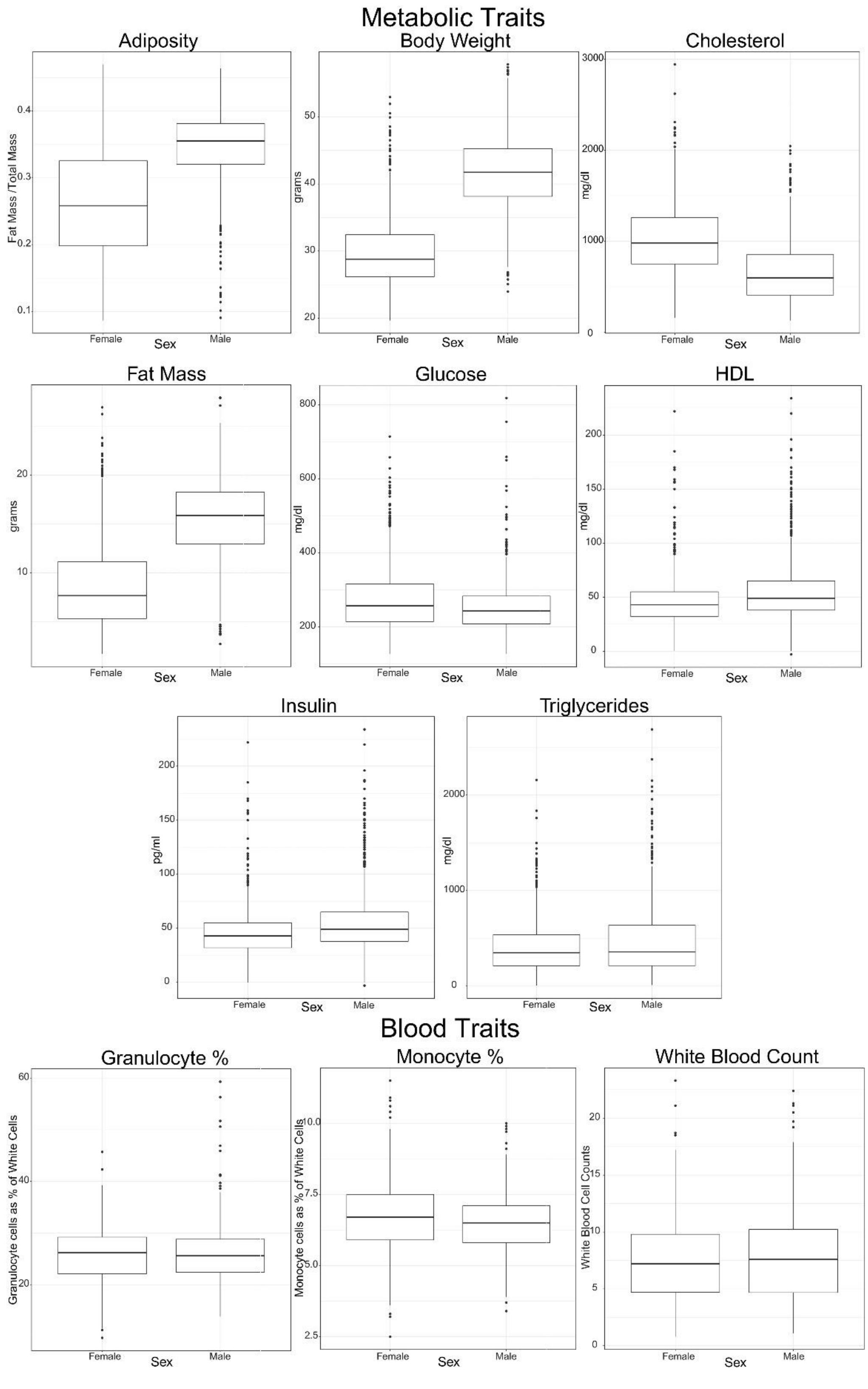
