## Supplementary material for "Concordance between male- and female-specific GWAS results helps define underlying genetic architecture of complex traits": Table S1

**Supplemental Table 1.** Genetic Correlation Between Metabolic Traits in the Hybrid Mouse Diversity Panel

**Males**

| | Body Weight | Cholesterol | Fat Mass | Glucose | HDL | Insulin | Triglycerides | Granulocyte % | Monocyte % | White Blood Count |
|---|---|---|---|---|---|---|---|---|---|---|
| Adiposity | 0.18 | -0.02 | 0.19 | 0.18 | 0.31 | -0.06 | -0.14 | -0.18 | 0.25 | -0.56 |
| Body Weight | | 0.48 | 0.86 | 0.19 | -0.03 | 0.63 | 0.6 | 0.39 | 0.52 | -0.46 |
| Cholesterol | | | 0.43 | 0.68 | -0.27 | 0.17 | 0.88 | 0.33 | 0.32 | -0.08 |
| Fat Mass | | | | 0.47 | 0.02 | 0.43 | 0.48 | 0.06 | 0.43 | -0.54 |
| Glucose | | | | | -0.23 | 0.17 | 0.56 | 0.11 | 0.28 | -0.5 |
| HDL | | | | | | -0.03 | -0.69 | -0.2 | -0.07 | 0.05 |
| Insulin | | | | | | | 0.34 | 0.15 | 0.15 | -0.6 |
| Triglycerides | | | | | | | | 0.48 | 0.47 | -0.12 |
| Granulocyte % | | | | | | | | | 0.83 | -0.41 |
| Monocyte % | | | | | | | | | | -0.48 |
| White Blood Count | | | | | | | | | | |

**Females**

| | Body Weight | Cholesterol | Fat Mass | Glucose | HDL | Insulin | Triglycerides | Granulocyte % | Monocyte % | White Blood Count |
|---|---|---|---|---|---|---|---|---|---|---|
| Adiposity | 0.18 | -0.08 | 0.1 | -0.26 | 0.14 | -0.04 | 0.13 | 0.05 | 0.22 | -0.38 |
| Body Weight | | 0.12 | 0.55 | 0.2 | 0.27 | 0.76 | 0.52 | 0.18 | 0.13 | -0.2 |
| Cholesterol | | | 0.21 | 0.52 | 0.03 | 0.19 | 0.65 | 0.26 | 0.23 | -0.07 |
| Fat Mass | | | | 0.32 | 0.25 | 0.83 | 0.57 | 0.11 | 0.21 | -0.35 |
| Glucose | | | | | -0.46 | 0.33 | 0.65 | 0.19 | 0.04 | -0.41 |
| HDL | | | | | | 0 | -0.42 | 0.13 | 0.08 | 0.2 |
| Insulin | | | | | | | 0.5 | 0.15 | 0.08 | -0.52 |
| Triglycerides | | | | | | | | 0.29 | 0.15 | -0.23 |
| Granulocyte % | | | | | | | | | 0.66 | -0.76 |
| Monocyte % | | | | | | | | | | -0.29 |
| White Blood Count | | | | | | | | | | |
