## Supplementary material for "Concordance between male- and female-specific GWAS results helps define underlying genetic architecture of complex traits": Table S2

**Supplemental Table 2.** Genetic Correlation of Traits Between Males and Females in the Hybrid Mouse Diversity Panel

| Trait | Genetic Correlation |
|---|---|
| Adiposity | 0.55 |
| Body Weight | 0.88 |
| Cholesterol | 0.41 |
| Fat Mass | 0.58 |
| Glucose | 0.86 |
| HDL | 0.17 |
| Insulin | 0.75 |
| Triglycerides | 0.58 |
| Granulocyte % | 0.59 |
| Monocyte % | 0.7 |
| White Blood Count | 0.35 |
