## Supplementary material for "Concordance between male- and female-specific GWAS results helps define underlying genetic architecture of complex traits": Table S3

**Supplemental Table 3.** Replicated Detection of Marginal Effects Using Less Stringent Threshold in Female Mice

| Trait | SNP | P-Value |
|---|---|---|
| Adiposity | JAX00692679 | 7.02E-06 |
| | JAX00692959 | 9.17E-06 |
| | JAX00691999 | 9.20E-06 |
| | JAX00692715 | 1.22E-05 |
| | JAX00692466 | 1.28E-05 |
| HDL | JAX00135190 | 8.12E-06 |
