## Supplementary material for "Concordance between male- and female-specific GWAS results helps define underlying genetic architecture of complex traits": Table S4

**Supplemental Table 4.** Bonferroni Significant Main Effects

Adiposity Marginal Effects in Male Mice

| SNP | Chr | ChrPos | PValue | Beta |
|---|---|---|---|---|
| JAX00076238 | 17 | 48009382 | 1.38E-13 | 2.35908629 |
| JAX00508560 | 2 | 157769293 | 7.77E-12 | -2.4303219 |
| JAX00313332 | 11 | 63005391 | 1.01E-11 | -3.2248415 |
| JAX00102330 | 2 | 160436398 | 2.04E-11 | -1.8373649 |
| JAX00125377 | 4 | 140666565 | 4.29E-11 | -3.7118865 |
| JAX00165119 | 8 | 96144288 | 5.62E-11 | 3.08113788 |
| JAX00377938 | 14 | 37266523 | 6.30E-11 | -2.2900286 |
| JAX00577770 | 5 | 30775108 | 1.02E-10 | -2.1851168 |
| JAX00670087 | 8 | 61204471 | 1.13E-10 | 2.99424997 |
| JAX00462778 | 18 | 66076404 | 1.45E-10 | -3.3149061 |
| JAX00277704 | 1 | 178288740 | 1.68E-10 | 1.51499564 |
| JAX00309417 | 11 | 44382107 | 2.72E-10 | 3.31720436 |
| JAX00387853 | 14 | 99805199 | 6.11E-10 | 3.03679907 |
| JAX00323401 | 11 | 119193315 | 2.20E-09 | -2.5510849 |
| JAX00439315 | 17 | 44401658 | 2.40E-09 | 2.8292101 |
| JAX00442095 | 17 | 54286241 | 2.96E-09 | 3.00011476 |
| JAX00569740 | 4 | 144791238 | 4.65E-09 | -2.6307678 |
| JAX00333580 | 12 | 52108546 | 5.42E-09 | -1.8651976 |
| JAX00359311 | 13 | 57278792 | 6.75E-09 | -2.3433622 |
| JAX00442337 | 17 | 55528851 | 7.54E-09 | 2.69198409 |
| JAX00351328 | 13 | 17697614 | 1.17E-08 | -2.7202128 |
| JAX00110698 | 3 | 96755927 | 1.58E-08 | -2.4812091 |
| JAX00164909 | 8 | 93326939 | 1.98E-08 | -3.3179359 |
| JAX00049996 | 14 | 10446252 | 2.11E-08 | -3.1031409 |
| JAX00440936 | 17 | 49593456 | 2.16E-08 | -2.4945998 |
| JAX00628084 | 6 | 138161899 | 2.59E-08 | 2.4392064 |
| JAX00039541 | 12 | 98629477 | 3.64E-08 | 3.17343983 |
| JAX00400446 | 15 | 50124649 | 4.72E-08 | 2.16789697 |
| JAX00294292 | 10 | 84542694 | 6.71E-08 | 2.30824459 |
| JAX00483267 | 2 | 10799042 | 7.14E-08 | 3.49252986 |
| JAX00060681 | 15 | 34428658 | 8.93E-08 | 2.31743556 |
| JAX00660982 | 8 | 12151377 | 9.03E-08 | -2.2177188 |
| JAX00384476 | 14 | 76113857 | 9.73E-08 | 2.22364933 |
| JAX00036628 | 12 | 59214598 | 1.07E-07 | 2.88419379 |

| | | | | |
|---|---|---|---|---|
| JAX00462088 | 18 | 64230628 | 1.30E-07 | 2.32281441 |
| JAX00303354 | 11 | 7179055 | 1.45E-07 | -2.0324507 |
| JAX00076511 | 17 | 51632056 | 1.48E-07 | 2.83520641 |
| JAX00504939 | 2 | 137599896 | 1.66E-07 | -1.9563215 |
| JAX00688378 | 9 | 28362190 | 1.66E-07 | -2.3795397 |
| JAX00127498 | 5 | 18327876 | 1.84E-07 | -2.3378667 |
| JAX00693290 | 9 | 47876230 | 1.98E-07 | -2.7477053 |
| JAX00246633 | 1 | 35405706 | 2.13E-07 | -2.2260967 |
| JAX00002346 | 1 | 34430864 | 2.19E-07 | 2.50208532 |
| JAX00629862 | 6 | 144634769 | 4.89E-07 | -2.5818042 |
| JAX00654103 | 7 | 123043169 | 5.08E-07 | -1.909699 |
| JAX00229764 | 6 | 68098210 | 5.28E-07 | -2.048248 |
| JAX00410644 | 15 | 95815634 | 5.58E-07 | -2.4344623 |
| JAX00430989 | 17 | 8924232 | 5.81E-07 | -2.874751 |
| JAX00466152 | 18 | 78349098 | 7.23E-07 | -2.2217261 |
| JAX00340033 | 12 | 83570083 | 7.85E-07 | 2.04292495 |
| JAX00603203 | 6 | 12636105 | 9.87E-07 | 1.94129793 |
| JAX00011772 | 1 | 160923234 | 1.06E-06 | 1.83715435 |
| JAX00397942 | 15 | 36110424 | 1.59E-06 | 2.16873117 |
| JAX00070786 | 16 | 69125405 | 1.68E-06 | 1.50430205 |
| JAX00318139 | 11 | 91502658 | 1.75E-06 | -2.0772628 |
| JAX00092911 | 2 | 33701416 | 1.83E-06 | 1.78255213 |
| JAX00632684 | 7 | 17808796 | 2.09E-06 | -2.2690794 |
| JAX00253170 | 1 | 63998987 | 2.54E-06 | -2.6538998 |
| JAX00682399 | 8 | 121786281 | 2.57E-06 | 2.27975099 |
| JAX00551981 | 4 | 57860723 | 2.63E-06 | -2.0904213 |

Body Weight Marginal Effects in Male Mice

| SNP | Chr | ChrPos | PValue | Beta |
|---|---|---|---|---|
| JAX00076238 | 17 | 48009382 | 1.27E-11 | 2.94073163 |
| JAX00165119 | 8 | 96144288 | 1.39E-09 | 3.55098095 |
| JAX00102330 | 2 | 160436398 | 5.34E-09 | -2.1014379 |
| JAX00313332 | 11 | 63005391 | 8.12E-09 | -3.5594638 |
| JAX00439315 | 17 | 44401658 | 1.69E-08 | 3.38967895 |
| JAX00277704 | 1 | 178288740 | 3.00E-08 | 1.68366226 |
| JAX00125377 | 4 | 140666565 | 7.72E-08 | -3.8212981 |
| JAX00462778 | 18 | 66076404 | 1.20E-07 | -3.405041 |
| JAX00508560 | 2 | 157769293 | 1.58E-07 | -2.4030478 |
| JAX00309417 | 11 | 44382107 | 1.65E-07 | 3.39416028 |

| | | | | |
|---|---|---|---|---|
| JAX00670087 | 8 | 61204471 | 1.75E-07 | 3.10349328 |
| JAX00387853 | 14 | 99805199 | 2.44E-07 | 3.18614453 |
| JAX00377938 | 14 | 37266523 | 2.70E-07 | -2.3994551 |
| JAX00049996 | 14 | 10446252 | 3.12E-07 | -3.5257515 |
| JAX00577770 | 5 | 30775108 | 3.39E-07 | -2.2964672 |
| JAX00039541 | 12 | 98629477 | 6.71E-07 | 3.6181323 |
| JAX00442337 | 17 | 55528851 | 9.94E-07 | 2.862735 |
| JAX00323401 | 11 | 119193315 | 1.54E-06 | -2.604417 |
| JAX00569740 | 4 | 144791238 | 1.82E-06 | -2.7382009 |
| JAX00092911 | 2 | 33701416 | 2.07E-06 | 2.12203371 |
| JAX00351328 | 13 | 17697614 | 2.19E-06 | -2.8196199 |
| JAX00688378 | 9 | 28362190 | 2.64E-06 | -2.6281694 |

Fat Mass Marginal Effects in Male Mice

| SNP | Chr | ChrPos | PValue | Beta |
|---|---|---|---|---|
| JAX00076238 | 17 | 48009382 | 1.38E-13 | 2.35908629 |
| JAX00508560 | 2 | 157769293 | 7.77E-12 | -2.4303219 |
| JAX00313332 | 11 | 63005391 | 1.01E-11 | -3.2248415 |
| JAX00102330 | 2 | 160436398 | 2.04E-11 | -1.8373649 |
| JAX00125377 | 4 | 140666565 | 4.29E-11 | -3.7118865 |
| JAX00165119 | 8 | 96144288 | 5.62E-11 | 3.08113788 |
| JAX00377938 | 14 | 37266523 | 6.30E-11 | -2.2900286 |
| JAX00577770 | 5 | 30775108 | 1.02E-10 | -2.1851168 |
| JAX00670087 | 8 | 61204471 | 1.13E-10 | 2.99424997 |
| JAX00462778 | 18 | 66076404 | 1.45E-10 | -3.3149061 |
| JAX00277704 | 1 | 178288740 | 1.68E-10 | 1.51499564 |
| JAX00309417 | 11 | 44382107 | 2.72E-10 | 3.31720436 |
| JAX00387853 | 14 | 99805199 | 6.11E-10 | 3.03679907 |
| JAX00323401 | 11 | 119193315 | 2.20E-09 | -2.5510849 |
| JAX00439315 | 17 | 44401658 | 2.40E-09 | 2.8292101 |
| JAX00442095 | 17 | 54286241 | 2.96E-09 | 3.00011476 |
| JAX00569740 | 4 | 144791238 | 4.65E-09 | -2.6307678 |
| JAX00333580 | 12 | 52108546 | 5.42E-09 | -1.8651976 |
| JAX00359311 | 13 | 57278792 | 6.75E-09 | -2.3433622 |
| JAX00442337 | 17 | 55528851 | 7.54E-09 | 2.69198409 |
| JAX00351328 | 13 | 17697614 | 1.17E-08 | -2.7202128 |
| JAX00110698 | 3 | 96755927 | 1.58E-08 | -2.4812091 |
| JAX00164909 | 8 | 93326939 | 1.98E-08 | -3.3179359 |
| JAX00049996 | 14 | 10446252 | 2.11E-08 | -3.1031409 |

| | | | | |
|---|---|---|---|---|
| JAX00440936 | 17 | 49593456 | 2.16E-08 | -2.4945998 |
| JAX00628084 | 6 | 138161899 | 2.59E-08 | 2.4392064 |
| JAX00039541 | 12 | 98629477 | 3.64E-08 | 3.17343983 |
| JAX00400446 | 15 | 50124649 | 4.72E-08 | 2.16789697 |
| JAX00294292 | 10 | 84542694 | 6.71E-08 | 2.30824459 |
| JAX00483267 | 2 | 10799042 | 7.14E-08 | 3.49252986 |
| JAX00060681 | 15 | 34428658 | 8.93E-08 | 2.31743556 |
| JAX00660982 | 8 | 12151377 | 9.03E-08 | -2.2177188 |
| JAX00384476 | 14 | 76113857 | 9.73E-08 | 2.22364933 |
| JAX00036628 | 12 | 59214598 | 1.07E-07 | 2.88419379 |
| JAX00462088 | 18 | 64230628 | 1.30E-07 | 2.32281441 |
| JAX00303354 | 11 | 7179055 | 1.45E-07 | -2.0324507 |
| JAX00076511 | 17 | 51632056 | 1.48E-07 | 2.83520641 |
| JAX00504939 | 2 | 137599896 | 1.66E-07 | -1.9563215 |
| JAX00688378 | 9 | 28362190 | 1.66E-07 | -2.3795397 |
| JAX00127498 | 5 | 18327876 | 1.84E-07 | -2.3378667 |
| JAX00693290 | 9 | 47876230 | 1.98E-07 | -2.7477053 |
| JAX00246633 | 1 | 35405706 | 2.13E-07 | -2.2260967 |
| JAX00002346 | 1 | 34430864 | 2.19E-07 | 2.50208532 |
| JAX00629862 | 6 | 144634769 | 4.89E-07 | -2.5818042 |
| JAX00654103 | 7 | 123043169 | 5.08E-07 | -1.909699 |
| JAX00229764 | 6 | 68098210 | 5.28E-07 | -2.048248 |
| JAX00410644 | 15 | 95815634 | 5.58E-07 | -2.4344623 |
| JAX00430989 | 17 | 8924232 | 5.81E-07 | -2.874751 |
| JAX00466152 | 18 | 78349098 | 7.23E-07 | -2.2217261 |
| JAX00340033 | 12 | 83570083 | 7.85E-07 | 2.04292495 |
| JAX00603203 | 6 | 12636105 | 9.87E-07 | 1.94129793 |
| JAX00011772 | 1 | 160923234 | 1.06E-06 | 1.83715435 |
| JAX00397942 | 15 | 36110424 | 1.59E-06 | 2.16873117 |
| JAX00070786 | 16 | 69125405 | 1.68E-06 | 1.50430205 |
| JAX00318139 | 11 | 91502658 | 1.75E-06 | -2.0772628 |
| JAX00092911 | 2 | 33701416 | 1.83E-06 | 1.78255213 |
| JAX00632684 | 7 | 17808796 | 2.09E-06 | -2.2690794 |
| JAX00253170 | 1 | 63998987 | 2.54E-06 | -2.6538998 |
| JAX00682399 | 8 | 121786281 | 2.57E-06 | 2.27975099 |
| JAX00551981 | 4 | 57860723 | 2.63E-06 | -2.0904213 |
