## Supplementary material for "Concordance between male- and female-specific GWAS results helps define underlying genetic architecture of complex traits": Table S5

**Supplemental Table 5.** Detection of Enrichment of Main and Interaction Effects, P-value < 0.05

| | Trait | Female Main Effects | Male Main Effects | Combined Main Effects |
|---|---|---|---|---|
| ***Expected Metabolic*** | | ***727*** | ***865*** | ***664*** |
| Metabolic | Adiposity | 1,214* | 988* | 1,459* |
| | Body Weight | 1,333* | 1,258* | 821* |
| | Cholesterol | 862* | 1,111* | 870* |
| | Fat Mass | 1,319* | 1,006* | 847* |
| | Glucose | 1,239* | 1,034* | 1,004* |
| | HDL | 633[&] | 1,047* | 919* |
| | Insulin | 1,522* | 1,166* | 925* |
| | Triglycerides | 1,116* | 1,052* | 951* |
| ***Expected Blood*** | | ***871*** | ***859*** | ***727*** |
| Blood | Granulocyte % | 1,093* | 900* | 1,206* |
| | Monocyte % | 1,134* | 954* | 1,221* |
| | White Blood Count | 1,070* | 1,040* | 1,166* |

*Significantly More Detected than Expected by Chance ($p < 0.05$)
[&]Significantly Fewer Detected than Expected by Chance ($p < 0.05$)
