## Supplementary material for "Concordance between male- and female-specific GWAS results helps define underlying genetic architecture of complex traits": Table S6

**Supplemental Table 6.** Number of Overlapping Significant Main Effects in Male Mice

| Trait | Number of Total Main Effect SNP | Number of Unique Main Effect SNPs | Number of Overlapping Main Effect SNPs |
|---|---|---|---|
| Adiposity | 58 | 4 | 54 |
| Body Weight | 22 | 0 | 22 |
| Fat Mass | 60 | 6 | 54 |

| Traits | Number of Overlapping Effects |
|---|---|
| Adiposity - Body Weight - Fat Mass | 20 |
| Only Adiposity - Body Weight | 1 |
| Only Adiposity - Fat Mass | 33 |
| Only Body Weight - Fat Mass | 1 |
