## Supplementary material for "Concordance between male- and female-specific GWAS results helps define underlying genetic architecture of complex traits": Table S7

**Supplemental Table 7.** Comparing Parks et al Significant Hits With HMDP Data

| Trait | Sex | Chr | Peak SNP | Position (Mb) | P-Value | Nearby Significant Loci (within 5 Mb) | R' |
|---|---|---|---|---|---|---|---|
| HOMA-IR | Male | 1 | rs32316569 | 175768939 | 5.22E-07 | JAX00277704 | 0.05 |
| Insulin | Male | 1 | rs31614030 | 173308737 | 4.66E-06 | NA | NA |
| Insulin | Female | 4 | rs27896920 | 71287837 | 4.46E-06 | NA | NA |
| Glucose | Male | 7 | rs3680765 | 49397123 | 1.52E-07 | NA | NA |
| Plasma Triglycerides | Male | 7 | rs31371723 | 76638409 | 4.64E-07 | NA | NA |
| Plasma Triglycerides | Female | 7 | rs33903345 | 65789990 | 7.94E-12 | NA | NA |
| Mesenteric Fat/Bodyweight | Male | 7 | rs32000744 | 67559538 | 2.71E-06 | NA | NA |
| Mesenteric Fat/Bodyweight | Female | 7 | rs32000744 | 67559538 | 8.73E-07 | NA | NA |
| HOMA-IR | Male | 9 | rs36804270 | 104875641 | 1.84E-06 | NA | NA |
| Glucose | Female | 11 | rs27001755 | 112944793 | 3.33E-07 | NA | NA |
| Insulin | Male | 15 | rs32269281 | 21154862 | 2.85E-06 | NA | NA |
| Mesenteric Fat/Bodyweight | Female | 15 | rs32263766 | 91948444 | 8.53E-07 | JAX00410644 | 0.02 |
| Mesenteric Fat/Bodyweight | Female | 15 | rs37298722 | 28527523 | 3.88E-06 | NA | NA |
| Plasma Triglycerides | Female | 17 | rs33060525 | 35595846 | 4.18E-06 | NA | NA |
| Mesenteric Fat/Bodyweight | Female | 17 | rs33115366 | 47452733 | 4.14E-06 | JAX00076238 | 0.07 |
