## Supplementary material for "Concordance between male- and female-specific GWAS results helps define underlying genetic architecture of complex traits": Table S8

**Supplemental Table 8.** Test of Normality for Metabolic and Blood Traits

| Trait | Female | Male |
|---|---|---|
| Adiposity | 2.45E-08 | <2.2E-16 |
| Body Weight | 4.18E-16 | 0.013 |
| Cholesterol | 7.33E-10 | <2.2E-16 |
| Fat Mass | <2.2E-16 | 7.30E-06 |
| Glucose | <2.2E-16 | <2.2E-16 |
| HDL | <2.2E-16 | <2.2E-16 |
| Insulin | <2.2E-16 | <2.2E-16 |
| Triglycerides | <2.2E-16 | <2.2E-16 |
| Granulocyte % | 0.22 | 3.89E-15 |
| Monocyte % | 0.016 | 0.11 |
| White Blood Count | 2.76E-09 | 8.77E-11 |
