## Supplementary material for "Concordance between male- and female-specific GWAS results helps define underlying genetic architecture of complex traits": Table S9

**Supplemental Table 9.** Main Effects of Trait Data Normalized or With Outliers Removed

Rank Based Normalized of Main Effects

| Trait | Female Bonferroni Significant Effects | Lambda | Male Bonferroni Significant Effects | Lambda | Combined Sex Bonferroni Significant Effects | Lambda |
|---|---|---|---|---|---|---|
| Adiposity | 0 | 1.09 | 58 | 0.99 | 1 (1) | 0.96 |
| Body Weight | 0 | 1.21 | 22 | 1.07 | 3 (3) | 0.58 |
| Cholesterol | 0 | 1.05 | 0 | 1.73 | 0 | 0.9 |
| Fat Mass | 0 | 1.13 | 60 | 1.53 | 0 | 0.76 |
| Glucose | 0 | 1.12 | 0 | 1.85 | 0 | 1.1 |
| HDL | 0 | 1.04 | 0 | 1.74 | 0 | 0.98 |
| Insulin | 0 | 1.19 | 0 | 1.76 | 0 | 0.93 |
| Triglycerides | 0 | 1.1 | 0 | 1.92 | 0 | 0.92 |
| Granulocyte % | 0 | 1.16 | 0 | 1.17 | 1 | 1.05 |
| Monocyte % | 0 | 1.17 | 0 | 1.07 | 0 | 1.13 |
| White Blood Count | 0 | 0.9 | 0 | 1.08 | 0 | 1.04 |

Box-Cox Based Normalized of Main Effects

| Trait | Female Bonferroni Significant Effects | Lambda | Male Bonferroni Significant Effects | Lambda | Combined Sex Bonferroni Significant Effects | Lambda |
|---|---|---|---|---|---|---|
| Adiposity | 0 | 1.14 | 63 (58) | 0.96 | 1 (1) | 0.94 |
| Body Weight | 0 | 1.1 | 22 (22) | 1.07 | 5 (5) | 0.54 |
| Cholesterol | 0 | 1.05 | 1 (0) | 1.04 | 0 | 0.91 |
| Fat Mass | 0 | 1.16 | 54 (54) | 0.98 | 3 (3) | 0.68 |
| Glucose | 0 | 1.13 | 0 | 1.15 | 0 | 1.1 |
| HDL | 0 | 1.15 | 2 (0) | 1.11 | 0 | 0.95 |
| Insulin | 0 | 1.19 | 0 | 1.07 | 0 | 1.03 |
| Triglycerides | 0 | 1 | 0 | 1.04 | 0 | 0.88 |
| Granulocyte % | 0 | 1.12 | 0 | 1.16 | 0 | 1.02 |
| Monocyte % | 0 | 1.07 | 0 | 1.17 | 0 | 1.13 |
| White Blood Count | 0 | 1.06 | 0 | 0.94 | 0 | 1.04 |

Main Effects Excluding Individual Outliers

| Trait | Female Bonferroni Significant Effects | Lambda | Male Bonferroni Significant Effects | Lambda | Combined Sex Bonferroni Significant Effects | Lambda |
|---|---|---|---|---|---|---|
| Adiposity | NA | NA | 35 (35) | 0.96 | NA | NA |
| Body Weight | 0 | 1.84 | 24 (22) | 1.65 | NA | NA |
| Cholesterol | 0 | 1.06 | NA | NA | 0 | 0.87 |
| Fat Mass | 0 | 1.87 | 55 (53) | 1.57 | NA | NA |
| Glucose | 0 | 1.76 | 0 | 1.18 | 0 | 1.08 |
| HDL | 0 | 0.16 | 0 | 1.09 | 0 | 1 |
| Insulin | 0 | 1.88 | 0 | 1.78 | 0 | 1 |
| Triglycerides | 1 | 1.62 | 0 | 1.83 | 0 | 0.95 |
| Granulocyte % | 0 | 1.17 | 0 | 1.11 | 0 | 1.03 |
| Monocyte % | NA | NA | 0 | 1.08 | 0 | 1.12 |
| White Blood Count | 0 | 0.91 | 0 | 1.06 | 0 | 1.06 |

Main Effects Excluding Strain Outliers

| Trait | Female Bonferroni Significant Effects | Lambda | Male Bonferroni Significant Effects | Lambda | Combined Sex Bonferroni Significant Effects | Lambda |
|---|---|---|---|---|---|---|
| Adiposity | NA | NA | NA | NA | NA | NA |
| Body Weight | 0 | 1.84 | NA | NA | NA | NA |
| Cholesterol | NA | NA | NA | NA | NA | NA |
| Fat Mass | NA | NA | NA | NA | NA | NA |
| Glucose | NA | NA | NA | NA | NA | NA |
| HDL | NA | NA | NA | NA | NA | NA |
| Insulin | NA | NA | 0 | 1.14 | 0 | 0.89 |
| Triglycerides | NA | NA | NA | NA | 0 | 0.92 |
| Granulocyte % | 0 | 1.17 | 0 | 1.14 | NA | NA |
| Monocyte % | NA | NA | NA | NA | NA | NA |
| White Blood Count | NA | NA | NA | NA | NA | NA |

*Parentheses notates the number of significant main effects replicated in the unadjusted analysis
