## Supplementary material for "Concordance between male- and female-specific GWAS results helps define underlying genetic architecture of complex traits": Table S10

**Supplemental Table 10.** Randomized Sex Main Effects in Adiposity

| | Female | Male |
|---|---|---|
| Permutation 1 | 2 | 0 |
| Permutation 2 | 0 | 4 |
| Permutation 3 | 0 | 0 |
| Permutation 4 | 0 | 1 |
| Permutation 5 | 17 | 0 |
| Permutation 6 | 0 | 0 |
| Permutation 7 | 2 | 2 |
| Permutation 8 | 0 | 3 |
| Permutation 9 | 11 | 4 |
| Permutation 10 | 1 | 3 |
| Permutation 11 | 0 | 0 |
| Permutation 12 | 0 | 2 |
| Permutation 13 | 0 | 0 |
| Permutation 14 | 0 | 0 |
| Permutation 15 | 0 | 0 |
| Permutation 16 | 0 | 0 |
| Permutation 17 | 0 | 1 |
| Permutation 18 | 0 | 2 |
| Permutation 19 | 0 | 0 |
| Permutation 20 | 0 | 0 |
| Permutation 21 | 0 | 10 |
| Permutation 22 | 0 | 0 |
| Permutation 23 | 0 | 1 |
| Permutation 24 | 0 | 0 |
| Permutation 25 | 0 | 0 |
| Permutation 26 | 0 | 0 |
| Permutation 27 | 0 | 0 |
| Permutation 28 | 0 | 10 |
| Permutation 29 | 0 | 0 |
| Permutation 30 | 0 | 0 |
| Permutation 31 | 0 | 0 |
| Permutation 32 | 0 | 0 |
| Permutation 33 | 1 | 0 |
| Permutation 34 | 0 | 0 |
| Permutation 35 | 0 | 0 |
| Permutation 36 | 0 | 0 |
| Permutation 37 | 0 | 0 |
| Permutation 38 | 0 | 0 |
| Permutation 39 | 0 | 0 |
| Permutation 40 | 0 | 0 |
| Permutation 41 | 0 | 0 |

| | | |
|---|---|---|
| Permutation 42 | 0 | 0 |
| Permutation 43 | 0 | 0 |
| Permutation 44 | 0 | 0 |
| Permutation 45 | 0 | 0 |
| Permutation 46 | 0 | 1 |
| Permutation 47 | 1 | 0 |
| Permutation 48 | 0 | 1 |
| Permutation 49 | 0 | 0 |
| Permutation 50 | 0 | 0 |
| Permutation 51 | 0 | 0 |
| Permutation 52 | 0 | 0 |
| Permutation 53 | 1 | 0 |
| Permutation 54 | 0 | 0 |
| Permutation 55 | 0 | 0 |
| Permutation 56 | 0 | 0 |
| Permutation 57 | 0 | 0 |
| Permutation 58 | 0 | 2 |
| Permutation 59 | 0 | 0 |
| Permutation 60 | 0 | 2 |
| Permutation 61 | 0 | 0 |
| Permutation 62 | 0 | 0 |
| Permutation 63 | 0 | 0 |
| Permutation 64 | 0 | 0 |
| Permutation 65 | 0 | 0 |
| Permutation 66 | 1 | 1 |
| Permutation 67 | 0 | 0 |
| Permutation 68 | 0 | 1 |
| Permutation 69 | 0 | 2 |
| Permutation 70 | 0 | 0 |
| Permutation 71 | 0 | 0 |
| Permutation 72 | 0 | 1 |
| Permutation 73 | 0 | 0 |
| Permutation 74 | 0 | 1 |
| Permutation 75 | 1 | 0 |
| Permutation 76 | 0 | 1 |
| Permutation 77 | 0 | 0 |
| Permutation 78 | 0 | 0 |
| Permutation 79 | 0 | 0 |
| Permutation 80 | 0 | 0 |
| Permutation 81 | 0 | 0 |
| Permutation 82 | 0 | 0 |
| Permutation 83 | 0 | 0 |
| Permutation 84 | 2 | 0 |

| | | |
|---|---|---|
| Permutation 85 | 0 | 0 |
| Permutation 86 | 0 | 13 |
| Permutation 87 | 0 | 0 |
| Permutation 88 | 0 | 0 |
| Permutation 89 | 0 | 0 |
| Permutation 90 | 0 | 0 |
| Permutation 91 | 0 | 0 |
| Permutation 92 | 0 | 0 |
| Permutation 93 | 0 | 4 |
| Permutation 94 | 0 | 2 |
| Permutation 95 | 0 | 0 |
| Permutation 96 | 0 | 0 |
| Permutation 97 | 0 | 1 |
| Permutation 98 | 0 | 0 |
| Permutation 99 | 0 | 3 |
| Permutation 100 | 1 | 0 |
