## Supplementary material for "Concordance between male- and female-specific GWAS results helps define underlying genetic architecture of complex traits": Table S11

**Supplemental Table 11.** Average Value and Direction of Betas for All Tested Main Effects

| Trait | Sex | Positive Beta | Negative Beta | Average Beta |
|---|---|---|---|---|
| Adiposity | Male | 35.29% | 65.71% | 0.04 |
| | Female | 25.71% | 74.29% | 0.004 |
| Body Weight | Male | 38.89% | 61.11% | 2.91 |
| | Female | 36.04% | 63.96% | 0.56 |
| Fat mass | Male | 40.00% | 60.00% | 2.45 |
| | Female | 31.11% | 68.89% | 0.3 |
