## Supplementary material for "Concordance between male- and female-specific GWAS results helps define underlying genetic architecture of complex traits": Table S12

**Supplemental Table 12.** Strains Used and Excluded from Metabolism and Blood Trait Analyses

| | Metabolism Traits* | | Blood Traits** | |
|---|---|---|---|---|
| **Strains** | **Male n** | **Female n** | **Male n** | **Female n** |
| 129X1/SvJ | 6 | 14 | 6 | 14 |
| A/J | 12 | 8 | 12 | 8 |
| AKR/J | 24 | 21 | 24 | 21 |
| AXB10/PgnJ | 4 | 2 | 4 | 2 |
| AXB12/PgnJ | 3 | 9 | 3 | 9 |
| AXB13/PgnJ | 10 | 12 | 9 | 11 |
| AXB15/PgnJ | 4 | 6 | 3 | 6 |
| AXB19/PgnJ | 13 | 6 | 13 | 6 |
| AXB19a/PgnJ | 14 | 12 | 14 | 12 |
| AXB19b/PgnJ | 4 | 4 | 4 | 4 |
| AXB2/PgnJ | 4 | 8 | 4 | 7 |
| AXB4/PgnJ | 3 | 0[&] | 3 | 0[&] |
| AXB5/PgnJ | 10 | 11 | 10 | 10 |
| AXB6/PgnJ | 3 | 3 | 3 | 3 |
| AXB8/PgnJ | 0[&] | 2 | 0[&] | 2 |
| BALB/cByJ | 12 | 4 | 12 | 4 |
| BALB/cJ | 16 | 18 | 16 | 17 |
| BTBR | 8 | 13 | 8 | 13 |
| BUB/BnJ | 10 | 19 | 10 | 19 |
| BXA1/PgnJ | 0[&] | 1 | 0[&] | 1 |
| BXA11/PgnJ | 7 | 7 | 7 | 7 |
| BXA14/PgnJ | 4 | 4 | 4 | 4 |
| BXA16/PgnJ | 4 | 3 | 4 | 3 |
| BXA2/PgnJ | 14 | 11 | 14 | 11 |
| BXA24/PgnJ | 18 | 13 | 18 | 13 |
| BXA25/PgnJ | 1 | 1 | 1 | 1 |
| BXA4/PgnJ | 4 | 4 | 4 | 4 |
| BXA7/PgnJ | 11 | 6 | 8 | 4 |
| BXA8/PgnJ | 10 | 9 | 10 | 9 |
| BXD1/TyJ | 1 | 5 | 1 | 5 |
| BXD11/TyJ | 2 | 4 | 2 | 4 |
| BXD12/TyJ | 2 | 0[&] | 2 | 0[&] |
| BXD13/TyJ | 4 | 3 | 4 | 3 |
| BXD14/TyJ | 4 | 3 | 4 | 3 |
| BXD18/TyJ | 0[&] | 3 | 0[&] | 3 |
| BXD2/TyJ | 4 | 1 | 4 | 1 |
| BXD20/TyJ | 4 | 6 | 4 | 6 |
| BXD21/TyJ | 11 | 7 | 11 | 6 |
| BXD22/TyJ | 2 | 1 | 2 | 1 |
| BXD24/TyJ-Cep290<rd16>/J | 0[&&] | 0[&&] | 0[&&] | 0[&&] |
| BXD28/TyJ | 2 | 0[&] | 2 | 0[&] |

| | | | | |
|---|---|---|---|---|
| BXD31/TyJ | 3 | 2 | 0& | 2 |
| BXD32/TyJ | 4 | 7 | 4 | 7 |
| BXD34/TyJ | 2 | 1 | 2 | 1 |
| BXD36/TyJ | 3 | 5 | 2 | 2 |
| BXD38/TyJ | 2 | 2 | 2 | 2 |
| BXD40/TyJ | 7 | 5 | 7 | 4 |
| BXD43/RwwJ | 5 | 4 | 5 | 4 |
| BXD44/RwwJ | 12 | 8 | 12 | 8 |
| BXD48/RwwJ | 8 | 18 | 8 | 18 |
| BXD5/TyJ | 3 | 6 | 2 | 6 |
| BXD50/RwwJ | 9 | 11 | 8 | 11 |
| BXD55/RwwJ | 9 | 17 | 9 | 17 |
| BXD56/RwwJ | 2 | 6 | 2 | 6 |
| BXD6/TyJ | 2 | 4 | 0& | 1 |
| BXD61/RwwJ | 14 | 5 | 12 | 5 |
| BXD64/RwwJ | 4 | 9 | 4 | 9 |
| BXD66/RwwJ | 7 | 3 | 7 | 3 |
| BXD68/RwwJ | 1 | 0& | 1 | 0& |
| BXD69/RwwJ | 3 | 12 | 3 | 12 |
| BXD70/RwwJ | 9 | 17 | 9 | 17 |
| BXD71/RwwJ | 2 | 4 | 2 | 4 |
| BXD73/RwwJ | 2 | 10 | 2 | 10 |
| BXD75/RwwJ | 12 | 10 | 12 | 8 |
| BXD79/RwwJ | 1 | 0& | 1 | 0& |
| BXD84/RwwJ | 8 | 6 | 8 | 6 |
| BXD86/RwwJ | 1 | 0& | 1 | 0& |
| BXD9/TyJ | 1 | 6 | 1 | 3 |
| BXH19/TyJ | 0& | 5 | 0& | 5 |
| BXH22/KccJ | 2 | 5 | 2 | 5 |
| BXH4/TyJ | 1 | 0& | 1 | 0& |
| BXH6/TyJ | 3 | 6 | 3 | 6 |
| BXH8/TyJ | 4 | 5 | 4 | 5 |
| BXH9/TyJ | 3 | 0& | 3 | 0& |
| C3H/HeJ | 12 | 13 | 12 | 13 |
| C57BL/6J | 15 | 16 | 15 | 16 |
| C57BLKS/J | 8 | 6 | 8 | 6 |
| C57L/J | 13 | 8 | 11 | 8 |
| C58/J | 5 | 4 | 4 | 4 |
| CBA/J | 8 | 5 | 7 | 5 |
| CE/J | 1 | 3 | 1 | 3 |
| CXB1/ByJ | 6 | 8 | 6 | 8 |
| CXB11/HiAJ | 1 | 1 | 1 | 1 |
| CXB12/HiAJ | 10 | 5 | 10 | 5 |
| CXB13/HiAJ | 8 | 9 | 8 | 9 |

| | | | | |
|---|---|---|---|---|
| CXB2/ByJ | 5 | 3 | 5 | 3 |
| CXB3/ByJ | 17 | 7 | 17 | 7 |
| CXB4/ByJ | 4 | 0[&] | 4 | 0[&] |
| CXB6/ByJ | 3 | 4 | 3 | 4 |
| CXB7/ByJ | 7 | 6 | 7 | 6 |
| CXB8/HiAJ | 2 | 5 | 0[&] | 1 |
| DBA/2J | 4 | 10 | 4 | 10 |
| FVB/NJ | 12 | 13 | 9 | 12 |
| I/LnJ | 10 | 6 | 10 | 6 |
| KK/HlJ | 3 | 5 | 3 | 5 |
| LG/J | 0[&] | 5 | 0[&] | 5 |
| LP/J | 2 | 5 | 2 | 5 |
| MA/MyJ | 4 | 4 | 4 | 4 |
| NOD/ShiLtJ | 5 | 6 | 5 | 6 |
| NON/ShiLtJ | 6 | 3 | 6 | 3 |
| NZB/BlNJ | 6 | 6 | 6 | 6 |
| NZW/LacJ | 11 | 12 | 11 | 12 |
| PL/J | 12 | 8 | 12 | 8 |
| RIIIS/J | 8 | 4 | 8 | 4 |
| SEA/GnJ | 11 | 7 | 11 | 7 |
| SJL/J | 10 | 7 | 10 | 7 |
| SM/J | 6 | 12 | 6 | 12 |
| SWR/J | 11 | 15 | 11 | 15 |

*Traits include Adiposity, Body Weight, Cholesterol, Fat Mass, Glucose, HDL, Insulin, [illegible]

**Traits include Granulocyte %, Monocyte %, and White Blo

[&]Removed due to missing phenotype data

[&&]Removed as strains model vision phenotypes
