## Supplementary material for "Concordance between male- and female-specific GWAS results helps define underlying genetic architecture of complex traits": Table S13

**Supplemental Table 13.** SNP Pruning and Haplotype Block Detection in the HMDP

| | **Original SNPs** | **LD Pruned** | **Haplotype Blocks** | **External SNPs** |
|---|---|---|---|---|
| Female Metabolic | 200,885 | 23,205 | 12,221 | 2,314 |
| Male Metabolic | 199,910 | 21,995 | 11,285 | 5,907 |
| Female Blood | 200,885 | 23,518 | 12,193 | 5,221 |
| Male Blood | 199,910 | 22,115 | 11,437 | 5,752 |
| Combined Metabolic and Blood | 195,190 | 22,284 | 12,229 | 5,219 |
